# Deconstructing the Cortical Sources of Frequency Following Responses to Speech: A Cross-species Approach

**DOI:** 10.1101/2021.05.17.444462

**Authors:** G. Nike Gnanateja, Kyle Rupp, Fernando Llanos, Madison Remick, Marianny Pernia, Srivatsun Sadagopan, Tobias Teichert, Taylor J. Abel, Bharath Chandrasekaran

## Abstract

Time-varying pitch is a vital cue for human speech perception. Neural processing of time-varying pitch has been extensively assayed using scalp-recorded frequency-following responses (FFRs), an electrophysiological signal thought to reflect integrated phase-locked neural ensemble activity from subcortical auditory areas. Emerging evidence increasingly points to a putative contribution of auditory cortical ensembles to the scalp-recorded FFRs. However, the properties of cortical FFRs and precise characterization of laminar sources are still unclear. Here we used direct human intracortical recordings as well as extra- and intracranial recordings from macaques and guinea pigs to characterize the properties of cortical sources of FFRs to time-varying pitch patterns. We found robust FFRs in the auditory cortex across all species. We leveraged representational similarity analysis as a translational bridge to characterize similarities between the human and animal models. Laminar recordings in animal models showed FFRs emerging primarily from the thalamorecepient layers of the auditory cortex. FFRs arising from these cortical sources significantly contributed to the scalp-recorded FFRs via volume conduction. Our research paves the way for a wide array of studies to investigate the role of cortical FFRs in auditory perception and plasticity.

**Significance Statement:** Frequency following responses (FFRs) to speech are scalp-recorded neural signals that inform the fidelity of sound encoding in the auditory system. FFRs, long believed to arise from brainstem and midbrain, have shaped our understanding of sub-cortical auditory processing and plasticity. Non-invasive studies have shown cortical contributions to the FFRs, however, this is still actively debated. Here we employed direct cortical recordings to trace the cortical contribution to the FFRs and characterize the properties of these cortical FFRs. With extra-cranial and intra-cranial recordings within the same subjects we show that cortical FFRs indeed contribute to the scalp-recorded FFRs, and their response properties differ from the sub-cortical FFRs. The findings provide strong evidence to revisit and reframe the FFR driven theories and models of sub-cortical auditory processing and plasticity with careful characterization of cortical and sub-cortical components in the scalp-recorded FFRs.

## INTRODUCTION

Time-varying pitch patterns are a vital component of all spoken languages. Periodicity, a critical cue for time-varying pitch (Plack et al., 2014) can be non-invasively assayed using the scalp-recorded frequency following responses (FFRs) in animals (Marsh et al., 1975; Smith et al., 1975; Ayala et al., 2017; Teichert et al., 2021) and humans (Coffey et al., 2019). Scalp-recorded FFRs reflect phase-locked activity from neural ensembles along the ascending auditory pathway (Worden and Marsh, 1968; Gerken et al., 1975; Gardi et al., 1979) and provide an integrative and non-invasive snapshot of pitch encoding in neurotypical and clinical populations (Chandrasekaran and Kraus, 2010b; Krizman and Kraus, 2019). In neurotypical populations, FFRs have been leveraged to demonstrate experience-dependent shaping of pitch patterns at pre-attentive stages of auditory processing. Periodicity encoding, as indexed by the FFRs, is found to be atypical in neurodevelopmental disorders (Abrams and Kraus, 2005; White-Schwoch et al., 2015), acquired neurological disorders (Kraus et al., 2016; Vander Werff and Rieger, 2017), and aging-related decline in auditory processing (Anderson et al., 2012; Bidelman et al., 2014; Presacco et al., 2016; Maruthy et al., 2017). Despite these critical contributions to our understanding of human auditory plasticity and the potential as an easy-to-record non-invasive biomarker, the neural sources of the scalp-recorded FFRs to pitch patterns are poorly understood.

For more than three decades, the inferior colliculus (IC) and the cochlear nucleus were considered the primary neural sources of the FFRs (Marsh et al., 1974; Smith et al., 1975; Yamada et al., 1980; Galbraith et al., 2000; Chandrasekaran and Kraus, 2010b; Gnanateja et al., 2012). Brainstem sources of scalp-recorded FFRs have been confirmed using cryogenic cooling of the IC (Smith et al., 1975) and resection of colliculo-cortical pathways (Greenberg et al., 1981) in animal models. Multichannel electroencephalography (EEG) recordings in humans also show that the subcortical sources predominantly contribute to the scalp-recorded FFRs (Galbraith et al., 2001; Bidelman, 2015, 2018; King et al., 2016). Furthermore, periodicity code of pitch is thought to be transformed into a rate or rate-place code in the upper brainstem (Plack et al., 2014). Recent studies using magnetoencephalography (MEG) and EEG have challenged these accounts and shown substantial cortical contributions to the scalp FFRs, with a distinct rightward cortical asymmetry (Coffey et al., 2016, 2021; Hartmann and Weisz, 2019; Gorina-Careta et al., 2021). However, non-invasive MEG and EEG studies require inferences based on distributed source modelling approaches which are relatively less sensitive to deep brain sources. FFRs to speech stimuli have also been demonstrated in the auditory cortex using direct intracortical recordings (Behroozmand et al., 2016; Guo et al., 2020). While converging evidence shows that FFRs can be recorded from the auditory cortex, it is not known if these FFRs are generated at the auditory cortex or if these are volume conducted electrical fields from the brainstem centers. The existing studies do not inform about the detailed arrangement of the current sources and sinks localized in the auditory cortex that can potentially give rise to these cortical FFRs. Additionally, it remains unclear how far the cortical FFRs are volume-conducted and contribute to the scalp-recorded FFRs.

Precise characterization of the laminar sources of FFR is challenging in human participants. Animal models that share anatomical and physiological similarity similarities to the human auditory pathway are invaluable in obtaining fine-grained information about the precise sources of the FFRs. In addition, animal models can also provide the freedom to record intracortical and scalp FFRs in the same animal to further deconstruct the cortical contribution to the scalp FFRs. The rhesus macaque and guinea pig are vocally communicating animals and have been successfully used as animal models to augment our understanding of the FFRs (Yamada et al., 1980; Chou et al., 2014; He et al., 2014; Ayala et al., 2017; Teichert et al., 2021). These animal models are highly similar to humans with respect to audible frequency range, auditory perceptual characteristics, and neuroanatomy (Sinnott et al., 1976; Sinnott and Kreiter, 1991; Kaas and Hackett, 2000; Rauschecker and Tian, 2000; Heffner and Heffner, 2007; Grimsley et al., 2012; Naert et al., 2019). Prior studies using these models have been restricted to the study of sub-cortical sources of FFRs (Yamada et al., 1980; He et al., 2014; Ayala et al., 2017). Considering the wide applicability of these animal models in understanding human auditory processing, we sought to track the cortical sources of FFRs in the two animal models and examine the extent to which they can aid in understanding FFRs in humans. Use of animal models to study FFRs aids in establishing a unified framework for studying the properties of FFRs. Such an approach can help in leveraging advanced species-specific scientific approaches that can provide different insights into the properties of FFRs in humans.

We used an integrative cross-species (human, rhesus macaque (Maq), and guinea pig (GP)) and cross-level (intra-cranial and extra-cranial) approach to deconstruct the cortical contribution to the FFR with unprecedented mechanistic detail. We examined human intracranial recordings with dynamic pitch varying stimuli (Mandarin tone stimuli) in two participants and confirmed the existence of cortical frequency following responses primarily localized to Heschl’s gyrus. We then applied representational similarity analysis (RSA) to demonstrate striking similarities in FFRs across species (Kriegeskorte et al., 2008; Barron et al., 2021), thereby establishing homologies across animal models to serve as a translational bridge. We further examined FFRs in the two animal models using fine spatial resolution to: a) deconstruct the laminar profile of the cortical sinks and sources of FFRs, and b) quantify cortical contribution to scalp FFRs using intracranial and extracranial recordings in the same animal using blind source separation and spectral profile estimates. Thus, by characterizing the FFRs with such unprecedented mechanistic detail using a cross-species approach, we demonstrate the existence of cortical sources of FFRs that emerge in thalamorecipient layers, and contribute to the scalp recorded FFRs.

## MATERIALS AND METHODS

### Stimuli

#### FFRs to Mandarin tones

The syllable /yi/ with four different pitch patterns (tones) were used to elicit the FFRs. These pitch patterns are linguistically-relevant in Mandarin and have been extensively used to examine experience-dependent auditory plasticity (Krishnan et al., 2010b, 2012; Lau et al., 2017, 2018; Llanos et al., 2017; Reetzke et al., 2018). The minimally contrastive F0 patterns are phonetically described as T1 (high-level, F0 = 129 Hz), T2 (low-rising, F0 ranging from 109 to 133 Hz), T3 (low-dipping, F0 ranging from 89 to 111 Hz), and T4 (high-falling, F0 ranging from 140 to 92 Hz). These stimuli were synthesized based on the F0 patterns (tones) derived from natural male speech production. All stimuli had a sampling rate of 48000 Hz and were 250 ms in duration. Stimuli were delivered using ER-3C insert earphones with the volume adjusted to a comfortable intensity level. The stimuli were presented in both condensation and rarefaction polarities to minimize potential contamination of the neural responses by the stimulus artifact and pre-neural cochlear microphonics (Skoe and Kraus, 2010a). A pseudorandom presentation was used where each stimulus had a ¼ probability of occurrence.

The overall number of stimuli sweeps per tone (T1-T4) differed across species, with *at least* 250 sweeps obtained in each species (Table 10).

### Intracranial and Scalp Electroencephalography in Humans

#### Participants

##### Human Participants for sEEG

FFRs were recorded intracranially in Hum1; a nine-year-old boy with drug-resistant epilepsy. The participant was right-handed, a native speaker of English, attending grade 4 in school. The participant underwent stereoelectroencephalography (sEEG) monitoring of the bilateral temporal lobes for localization of his seizure focus. An opportunity to record sEEG from both temporal lobes to study bilateral auditory processing in the same participant is unique. This participant had no other relevant medical history. To assess generalizability, we also recorded intracranial FFRs in a second participant Hum2; a sixteen-year-old boy with drug-resistant epilepsy. The participant was right-handed, was a native speaker of English, and had completed grade 9 in school. The participant underwent stereoelectroencephalography (sEEG) monitoring of broad right frontotemporal regions for localization of his seizure foci. In both participants, the choice of electrode insertion was based purely on clinical necessity for evaluation of focal epilepsy. Both participants’ family gave written informed consent to participate in the study. All research protocols were approved by the Institutional Review Board of the University of Pittsburgh.

##### Human Participants for EEG

Data from a previously published study (Reetzke et al., 2018) with twenty participants in the age range of 18 to 24 years (12 females) was reanalyzed in this study. All the participants were monolingual native speakers of English. All the participants had hearing sensitivity within 20 dB HL across octave frequencies from 250 to 8000 Hz. Written informed consent was obtained from the participants before inclusion in the study. The research protocols used were approved by the Institutional Review Board of the University of Texas, Austin.

#### Electrophysiological recordings in Humans

##### Stereoelectroencephalography in Humans

Stereoelectroencephalography (sEEG) electrodes were inserted into the brain using robot-assisted implantation (Abel et al., 2018; Faraji et al., 2020). Twenty electrode trajectories in Hum1 (129 active electrode contacts) and eighteen trajectories Hum2 (226 active electrode contacts) were inserted along different brain regions to test seizure localization hypotheses based on non-invasive evaluations (Chabardes et al., 2018). Each electrode had between 8-12 cylindrical contacts with a length of 2mm and a diameter of 0.8mm. The distance between each electrode contact was 3.5mm. The choice of electrode sampling (spatial resolution) across the trajectories was made based on clincal necessity. Anatomical locations of the electrode sites were obtained using high-resolution CT and structural MRI. The electrode locations from the CT scan were coregistered with the structural MRI to precisely locate the anatomical locations of each electrode. A cortical reconstruction was generated from the MRI using Freesurfer (Fischl, 2012), and electrodes were localized using CT coregistration in Brainstorm (Tadel et al., 2011). The sEEG signals were recorded with a Grapevine Nomad processor (Ripple LLC, Salt Lake City, UT) and the accompanying Trellis recording software. The sEEG was recorded at a sampling rate of 1000 Hz and an online notch-filter was applied at 60/120/180 Hz to reduce electrical line interferences. The audio signal was synchronously recorded by the Grapevine system at a sampling rate of 30000 Hz. The auxiliary audio channel was used to mark the onset times of each stimulus in the sEEG recordings. The participants passively listened to the Mandarin vowels. Neither participant had seizure foci in the temporal lobe, nor had any active seizure activity in the during the experiment.

##### Scalp Electroencephalography in Humans

To understand how the FFRs recorded at the cortex compare with the FFRs that are conventionally obtained from the scalp, we utilized scalp-recorded FFR dataset from a previously published study. This dataset also was used to establish a translation bridge between scalp-recorded FFRs and cortical FFRs across humans and animal models. The details of scalp EEG are provided briefly here, complete details are available in the source paper (Reetzke et al., 2018). Scalp-recorded FFRs were recorded from the 20 human participants (10 female) using EEG. EEG was recorded with a single AgCl electrode placed on the scalp that was referenced to the left mastoid, and the ground was placed on the opposite mastoid. Brainvision EEG system was used to record the EEG activity. A dedicated preamplifier (EP-preamp) connected to the actichamp amplifier with a gain setting of 50x.

### Intracranial and scalp electroencephalography in Rhesus Macaque

#### Subjects

The EEG experiments and intracortical recordings were performed on two adult male macaque monkeys (Macaca mulatta, Maq1, Maq2). The treatment of the monkeys was in accordance with the guidelines set by the U.S. Department of Health and Human Services (National Institutes of Health) for the care and use of laboratory animals. All methods were approved by the Institutional Animal Care and Use Committee at the University of Pittsburgh. The animals were between 5 and 11 years old and weighed between 8-11 kg at the time of the experiments.

#### Cranial EEG recordings

Details of the cranial EEG recordings have been reported previously (Teichert, 2016; Teichert et al., 2016). Briefly, EEG electrodes manufactured in-house from medical grade stainless steel were implanted in 1mm deep, non-penetrating holes in the cranium of Maq1. All electrodes were connected to a 36-channel Omnetics connector embedded in dental acrylic at the back of the skull. The 33 electrodes formed regularly-spaced grids covering roughly the same anatomy covered by the international 10-20 system (Li and Teichert, 2020). All the electrodes were referenced to an electrode placed at Oz.

#### Intracranial recordings in primary auditory cortex

For the single-tipped sharp electrode recordings in Maq1, neural activity was recorded with a chronically implanted 96 channel electrode array with individually movable electrodes (SC96 from Graymatter). For the laminar recording in Maq2, neural activity was recorded with a 24 channel laminar electrode (S-Probe from Plexon) positioned approximately perpendicular to the left superior temporal plane’s orientation. The depth of the probe was adjusted iteratively until the prominent sound-evoked supra-granular source was located slightly above the center of the probe. At the time of the experiments, 12 of the electrodes were positioned in or close enough to the superficial layers of the auditory cortex to pick up frequency-tuned local field potentials. Six of these electrodes also picked up frequency-tuned multi-unit activity, suggesting that they were located in layer III or below. The devices in both animals were implanted over the right hemisphere in a way that allowed electrodes to approach the superior temporal plane approximately perpendicular.

#### Experimental Setup

All experiments were performed in small (4’ wide by 4’ deep by 8’ high) sound-attenuating and electrically insulated recording booths (Eckel Noise Control Technology). Animals were positioned and head-fixed in custom-made primate chairs (Scientific Design). Neural signals were recorded with a 256-channel digital amplifier system (RHD2000, Intan) at a sampling rate of 30 kHz.

Experimental control was handled by a Windows PC running an in-house modified version of the Matlab software-package monkeylogic. Sound files were generated prior to the experiments and presented by a sub-routine of the Matlab package Psychtoolbox. The sound-files were presented using the right audio-channel of a high-definition stereo PCI sound card (M-192 from M-Audiophile) operating at a sampling rate of 96 kHz and 24 bit resolution. The analog audio-signal was then amplified by a 300 Watt amplifier (QSC GX3). The amplified electric signals were converted to sound waves using a single element 4 inch full-range driver speaker (Tang Band W4-1879) located 8 inches in front of the animal, and presented at an intensity of 78 dB SPL. To determine sound onset with high accuracy, a trigger signal was routed through the unused left audio channel of the sound card directly to one of the analog inputs of the recording system. The trigger pulse was stored in the same stereo sound-file and was presented using the same function call. Hence, any delay in the presentation of the tone also leads to an identical delay in the presentation of the trigger. Thus, sound onset could be determined at a level of accuracy that was limited only by the sampling frequency of the recording device (30kHz: corresponding to 33 μsec).

### Cranial electroencephalography and intracranial recordings in guinea pigs

#### Subjects

The cranial EEG and intracranial recordings were performed on two wild-type (∼ 8 months old), pigmented guinea pigs (GPs, GP1 and Gp2; Cavia porcellus; Elm Hill Labs, Chelmsford, MA), weighing ∼600 – 800g. All experimental procedures were conducted according to NIH Guidelines for the care and use of laboratory animals and were approved by the Institutional Animal Care and Use Committee (IACUC) of the University of Pittsburgh.

Prior to commencing recordings, a custom headpost for head fixation, skull screws that served as EEG recording electrodes or reference electrodes for intracranial recordings, and recording chambers for intracranial recordings were surgically implanted onto the skull using dental acrylic (C & B Metabond, Parkell Inc.) following aseptic techniques under isoflurane anesthesia. Analgesics were provided for three days after surgery, and animals were allowed to recover for ∼10 days. Following recovery, animals were gradually adapted to the recording setup and head fixation for increasing durations of time.

#### Experimental Setup

All recordings were performed in a sound-attenuated booth (IAC) whose walls were covered with anechoic foam (Pinta Acoustics). Animals were head-fixed in a custom acrylic enclosure affixed to a vibration isolation tabletop. Stimuli were presented using Matlab (Mathworks, Inc). Digital stimulus files sampled at 100 kHz were converted to an analog audio signal (National Instruments, USA), attenuated (Tucker-Davis Technologies, USA), power-amplified (Tucker-Davis Technologies, USA) and delivered through a calibrated speaker (4” full-range driver, TangBand, Taiwan) located ∼0.9 m in front of the animal. Stimuli were presented at ∼75 dB SPL.

#### Cranial EEG recordings

FFRs were acquired from unanesthetized, head-fixed, passively-listening GPs using a vertical electrode montage. Scalp-recorded activity was collected via a stainless-steel skull screw (Fine Science Tools, USA). Reference and ground conductive adhesive hydrogel electrodes (Foam electrodes, Kendall™ or Covidien Medi-trace®, USA) were placed on the high forehead and mastoid respectively. Signals were acquired using a multichannel neural processor (Ripple Inc., USA).

#### Intracranial recordings in primary auditory cortex

Intracortical recordings were performed in unanesthetized, head fixed, passively-listening GPs. Small craniotomies (1 – 2 mm. Diameter) were performed within the implanted recording chambers, over the expected anatomical location of primary auditory cortex. Neural activity was recorded using high-density 64-channel multi-site electrode (Cambridge Neurotech), inserted roughly perpendicular to the cortical surface. The tip of the probe was slowly inserted to a depth of ∼2 mm. After a brief waiting period to allow the tissue to settle, signals were acquired using a multichannel neural processor (Ripple Scout).

A summary of the participant information, stimulus and acquisition parameters across the three species are provided in Table 1.

**Table 1:** Details of participants, stimulus and recording parameters

| Parameters | Human sEEG | Human EEG | Macaque intracranial | Macaque extracranial | Guinea pig Intracranial | Guinea pig Extracranial |
| --- | --- | --- | --- | --- | --- | --- |
| Number of Participants | 2 | 20 | 2 | 2 | 2 | 2 |
| Electrode location | Intracranial electrode with cylindrical contacts.<br>*Hum1*: 129 electrode contacts in Bilateral temporal lobes<br>*Hum2*: 229 electrode contacts in the right hemisphere;temporal, frontal, and parietal lobes | Surface electrodes on the Vertex and Mastoids | Maq1: 96 Sharp electrodes on the surface of the auditory auditory and motor cortices.<br>Maq2: 24 laminar probe electrodes the layers of the auditor cortex | Maq1: 33 EEG electrodes . | Laminar probe in the auditory cortex | Surface electrodes at Cz and T4. |
| Transducer | ER-3C | ER-3C | Sound-field speaker | Sound-field speaker | Sound-field Speaker | Sound-field Speaker |
| Intensity | Comfortable level | 75 dBSPL | 78 dBSPL | 78 dBSPL | 75 dBSPL | 75 dBSPL |
| Number of sweeps | 800 | 1000 | 1000 | 1000 | 250 | 2000 in GP1<br>250 in GP2 |
| Interstimulus interval | 60 ms | 128-168 ms | 250 ms | 250 ms | 250 ms | 100 ms |

### Data Processing and Analyses

#### FFR Preprocessing and Analysis for sEEG in humans

A linear regression-based method was used to remove the harmonics of the power line interference from the data, using the cleanline plugin in EEGLAB (Delorme and Makeig, 2004). The raw sEEG was high pass filtered using a third-order zero-phase shift butterworth filter. No low pass filter was used, as the sampling rate was 1000 Hz resulting in an effective low pass frequency of 500 Hz. Time-locked sEEG epochs were extracted for all vowels in both polarities. The epochs that exceeded amplitudes of 75 µV were rejected. The FFRs in both the polarities for each vowel were averaged to obtain a total of four FFR waveforms (one for each vowel). Four FFR waveforms were obtained for each electrode.

The inter-trial phase locking value (ITPC) was estimated to assess the frequencies at which the cortical units phase-lock. The single-trial FFRs were decomposed into a spectrogram representation using time-frequency analysis timefreq.m in EEGLAB. Specifically, 130 wavelets between 70 and 200 Hz with equal widths were used for the time-frequency decomposition. The complex valued time-frequency vectors were divided by their magnitude to obtain unit vectors at every frequency and time point. These unit vectors in the time-frequency domain were averaged across trials to obtain resultant vectors. The absolute magnitude of the resultant vectors at each time and frequency point was used to obtain the ITPC spectrogram (Figure 1). ITPCs ranged from 0 to 1, with 0 indicating no phase-locking and 1 indicating perfect phase-locking across trials. These ITPCs provide information about the extent of phase-locking at different frequencies and latencies without the confounds of differences in absolute FFR magnitude.

**Figure 1.**
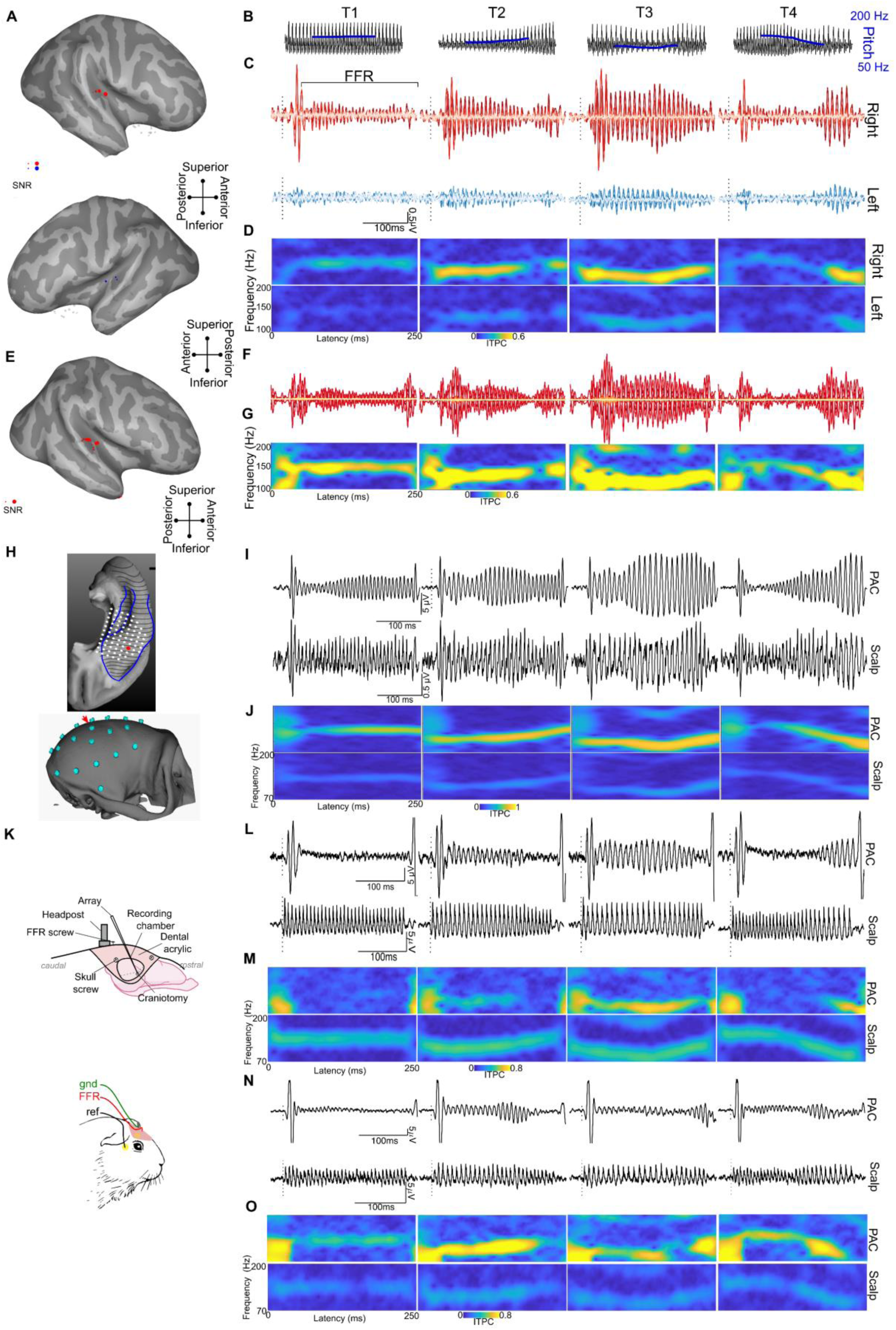
Cross-species and cross-level approach to characterizing the functional properties of cortical FFRs. **A-G** human data, **H-J** Macaque Data, **K-O** Guinea pig data **A** and **E** show the Location of the sEEG electrodes (projected to the surface for visualization) projected on the inflated brain surfaces of the human participants Hum1 and Hum2 respectively. Electrodes in gray do not show any significant FFRs above the pre-stimulus baseline (p<0.01 on paired t-tests on bootstrapped samples). Electrodes marked in red or blue show significant FFR magnitudes above the pre-stimulus baseline, where the size of the marker is proportional to the signal to noise ratio for FFRs. **B** Waveforms of the Mandarin tones /yi/; T1 (high-level fundamental frequency (F0)), T2 (low-rising F0), T3 (low-dipping F0), and T4 (high-falling F0). **C** & **F** FFR waveforms from sEEG electrodes in both right (top panel – right) and left (bottom panel - blue) temporal lobes in in Hum1 and Hum2 respectively. **D** & **G** FFR inter-trial phase coherence (ITPC) at all time points and frequencies with a spectrogram for the sEEG FFRs in Hum1 and Hum2 respectively. The ITPC spectrograms shown are only for the electrodes with the highest FFR amplitude in each hemisphere (located on the HG). **H** *Top* Locations of the semi-chronic sharp electrodes (white dots) used to record FFRs in the primary auditory cortex (PAC) of the macaque. FFRs from a representative example electrode highlighted in red. *Bottom* Layout of the EEG Electrode grid. FFR responses analyzed here were from the electrode marked with the red arrow. **I** Waveforms show simultaneously recorded FFRs from macaque primary auditory cortex (Top - PAC) and skull (Bottom - ∼ Cz). **J** ITPC spectrograms of FFRs in the macaque at both PAC and scalp. **K** intracranial and extracranial electrode setup in the guinea pigs GP1 and GP2. **L** & **N** show FFR waveforms to the four Mandarin tones recorded from the best laminar depth electrode placed in the PAC (*top*) and from a surface scalp electrode (*bottom*) of the guinea pigs GP1 and GP2 respectively. **M** & **O** show ITPC spectrograms of FFRs in guinea pigs GP1 and Gp2.

#### FFR processing and Analysis for EEG in Humans

The EEG data from the humans was bandpass filtered from 80 to 1000 Hz and epoched from -25 to 250 ms (re:stimulus onset). Baseline correction was applied on each epoch, and the epochs exceeding an amplitude of +/-35 µV were excluded from further analysis.

#### FFRs to Mandarin tones in macaques and guinea pigs

The raw data were high pass filtered using a second-order zero-phase shift FIR filter. Time-locked epochs were extracted for all vowels in both polarities. The epochs that exceeded amplitudes of 250 µV were rejected. The FFRs in both the polarities for each tone were averaged to obtain a total of four FFR waveforms (one for each tone).

Custom Matlab and R routines were used to filter and average signals appropriately to obtain local field potentials (LFPs) and multi-unit activity (MUA) from the macaque and GP recordings. The current source density (CSD) in the macaque was computed from the LFP signals derived from the electrodes with 150 µm spacing, spatially smoothing the LFP signal using a Gaussian filter (SD = 250 µm), and obtaining the second spatial derivative method using the finite difference approximation. In the GP, CSD was computed from the LFP signals derived from alternate electrode contacts (60 µm spacing), spatially smoothing the LFP signal using a Gaussian filter (SD = 125 µm) and obtaining the second spatial method using the finite difference approximation. The sink with the earliest latency in the CSD, post-stimulus onset, was used to identify the thalamorecipient layers.

#### Representational similarity analysis (RSA) of FFRs to Mandarin tones

RSA was used to establish homologies between the species and assess similarities across scalp and cortical FFRs. RSA was performed on the accuracies of a machine learning model to decode the four mandarin tones (pitch patterns) based on the FFRs. A hidden Markov model (HMM) classifier was used as the machine learning model and was trained to decode the Mandarin tones based on the FFR pitch tracks (Llanos et al., 2017). A detailed description of the HMM-based decoding approach can be found in a previously published methods paper (Llanos et al., 2017). The averaging size of the HMM was dynamically adjusted to obtain equivalent classification accuracies across the different levels (scalp, cortex) and species. The averaging sizes used were 150 trials for human scalp FFR, 24 and 2 trials for sEEG FFR in Hum1 and Hum2 respectively, 150 trials for macaque scalp FFR, 4 trials for macaque PAC FFR, 6 trials for the guinea pig scalp FFRs, and 16 trials for the GP PAC FFRs. The confusion matrices of decoding patterns (proportion correct) were extracted and used for further representational similarity analysis. Multi-dimensional scaling analysis was performed on the confusion matrices (diagonals removed) to assess if the decoding patterns across levels and species are similar. Procrustes analysis was performed to rotate and transform all the MDS representation to the same scale to facilitate comparison across species and levels. A similarity matrix was derived from the confusion matrices (diagonals removed) across-levels and-species using Pearson’s product-moment correlations and the significance of these correlations were also assessed.

#### Comparison of FFRs recorded at the scalp and the auditory cortex

##### Spectro-temporal Measures

In the animal models, we had the opportunity to recorded FFRs from the scalp and the cortex in the same animal (Maq1, GP1, and GP2). We compared the FFR power spectra at the scalp and cortex to analyze similarities and differences in the spectral composition between the scalp and cortical FFRs. Welch’s power spectral density (PSD) estimates of the FFRs were obtained with a 1024 point hamming window with 50% overlap, to obtain a smoothed spectral estimate of the FFRs. The PSDs of all the four tones were averaged to obtain an average spectral composition of the FFRs. The PSD of the scalp and cortical FFRs were normalized by setting their maximum magnitude to 0 dB. This facilitated the comparison of the spectral decay and relative differences in encoding of the F0 and the higher harmonics at the scalp and the cortex. Due to the high amplitude of the FFR at the F0, the normalization process essentially normalized the F0 magnitude, which facilitated the inspection of decay in magnitude of the high frequency components in the FFRs with reference to the F0 magnitude.

Due to time-varying F0 trajectories in the stimuli, it is challenging to infer the FFR temporal properties from a singular estimate of cross-correlation latency based on the raw FFR waveforms. This is especially challenging when contributions of different underlying sources of FFRs are also expected to change depending on stimulus frequency. Thus, we compared the latencies between the scalp and cortical FFRs in the time-frequency domain. We decomposed the FFRs into a time-frequency representation using the continuous wavelet transformation using the Morse wavelet implementation in the Matlab wavelet analysis toolbox. The real-valued wavelet-decomposed waveforms at every frequency were then cross-correlated between the scalp and cortical FFRs. The cross-correlation lags with the highest absolute correlation coefficient were estimated. This analysis provided a latency estimate (cross-correlation lag) and magnitude of similarity (Pearson’s *r*) for every frequency in the FFRs. The latencies measured here are with the scalp FFRs as reference. Thus, positive latencies indicate that the cortical FFRs lag scalp FFRs in latency. One scalp and one intracortical electrode with the best signal-noise ratio were chosen for this analysis. In Maq1, scalp and intracortical FFRs were obtained simultaneously. In GP1, scalp and intracortical FFRs were obtained in two separate sessions, while in GP2 both scalp and intracortical FFRs were obtained simultaneously. The FFRs from the cortex were derived from the electrode with the maximum signal-to-noise ratio in laminar FFR-LFP recordings.

A cross-spectral power density estimate was also obtained to assess the similarity in power between the scalp and cortical FFRs irrespective of the latency difference. This estimate is useful in getting an objective metric of similarity in spectral properties of the FFRs at the cortex and the scalp, irrespective of the differences in temporal properties. The Cross-spectral densities were obtained using Welch’s periodogram method with 50% overlapping 1024-point hamming windows. The absolute power of the cross-spectral densities was averaged across the FFRs for the four Mandarin tones and overlaid on the plot of frequency-wise latency comparison to obtain a unified inference of the spectro-temporal similarities in the FFRs at the scalp and cortex.

The above measures show differences and similarities in the scalp and cortically recorded FFRs. Multiple volume-conducted fields in the cortex and sub-cortex that lead to constructive and destructive interferences drive these differences and similarities, and the above measures may not be sensitive to differentiate these fields. Thus, we further used a blind source separation approaches using independent component analysis that can separate spectro-temporally overlapping components arising from different neural sources (Makeig et al., 2004). ICA extracts mutually independent components disentangling the superposed electrical fields from the different electrical dipoles across the auditory cortical subcortical regions and their projection to the scalp and the cortical electrodes. The electrodes placed at the cortex though record activity from the cortical ensembles in their proximity, they also pick up electrical fields from the subcortical regions and the surrounding auditory cortical fields. With ICA, we can decompose the neural sources underlying the electrical activity across the cortical and scalp electrodes and trace the differential projection of neural sources to the individual electrodes.

ICA was applied on 33 scalp electrodes and 96 sharp electrodes from the cortical surface (spanning the surface of superior temporal plane and motor cortices) in the macaque, and 25 laminar depth electrodes in the auditory cortex and two scalp electrodes (Cz and T4) in the guinea pig. The averaged FFRs corresponding to all four Mandarin tones were input into the ICA decomposition. The Infomax algorithm (runica.m) in EEGLAB toolbox (Delorme and Makeig, 2004) was used for ICA decomposition of the neural data. A principal component analysis was performed to reduce the dimensionality of the signal and restricted the decomposition to components that explained 96% (Macaque) and 99% (Guinea Pig) of the original variance in the data. The percentage variance accounted for by each independent component (ICA) was estimated (eeg_pvaf.m). The ICAs that each explained greater than 10% of the variance in the FFRs were retained for further analysis. The spatial weights of the ICAs were derived as the pseudoinverse of the product of the ICA weights and the ICA sphering matrix. These spatial weights were used to assess the spatial layout of the volume-conducted propagation of the different ICAs. The PVAFs of the ICAs at each scalp and cortical electrode were estimated to obtain the contribution of the ICAs to each of the electrodes. The emphasis of this analysis was to assess the percent contribution of the cortical ICAs to the scalp-recorded FFRs. The latencies of the ICAs were assessed by cross-correlation with the stimulus waveform. These latencies were also used to infer the potential generators of the ICAs, with earlier latencies corresponding to more subcortical sources. The power coherence was estimated between each of these ICAs and the stimulus waveform using cross-power spectral density (cpsd.m). This analysis aided in inferring the differential pattern in the decline of power coherence across the cortical ICAs and subcortical ICAs. The power coherence metric is not a measure of phase-locking but just the power coherence between the stimulus and the ICAs.

**Code availability:** The analysis codes will be provided to the readers on request.

## RESULTS

### FFRs in the human auditory cortex to vowels with time-varying pitch contours

FFRs were recorded from stereotactically implanted electrodes (Fig. 1A and 1E) in two participants while they listened to the pitch varying Mandarin vowels. In both participants, the location of the electrodes was based on clinical necessity. In Hum1, the electrodes were implanted in both hemispheres with electrodes spanning across the superior temporal plane, superior temporal gyri/sulci, middle temporal gyri/sulci, and insula. In Hum2 the electrodes were implanted only in the right hemisphere spanning the frontal, parietal, and temporal lobes, the superior temporal plane, and the insula.

We analyzed time- and phase-locked neural activity to the periodicities in the stimulus. Robust FFRs (Fig 1C & 1F) with amplitudes above pre-stimulus baseline (*p*<0.05 on permutations-based t-tests between pre-stimulus baseline and FFRs on bootstrapped FFR trials) were observed in the electrode contacts in the Heschl’s gyrus (HG) and the Planum Temporale in both subjects (6/129 electrode contacts in Hum1, and 10/226 electrode contacts in Hum2) (Fig 1A & 2E). Electrode contacts farther from HG did not show FFR like responses that were significantly above the pre-stimulus baseline level (*p*<0.05 on permutations-based t-tests between pre-stimulus baseline and FFRs on bootstrapped FFR trials). Thus, further FFR analyses were restricted to the electrodes along the HG.

We used four-pitch variants of the vowel /yi/ (Fig 1B), referred to as Mandarin tones to elicit the FFRs (Fig 1C). These Mandarin tones have been extensively used to record FFRs to study the neurophysiology of pitch processing and associated plasticity in humans (Krishnan et al., 2010a; Llanos et al., 2017; Lau et al., 2018; Reetzke et al., 2018). The Mandarin tones are phonetically described as; T1 (high-level F0), T2 (low-rising F0), T3 (low-dipping F0), and T4 (high-falling F0). Morphologically, the time-locked averaged sEEG responses to the Mandarin tones showed robust onset responses followed by FFRs that lasted throughout the stimulus duration. We refer to the FFRs recorded from electrode contacts in close proximity to or directly within the auditory cortex as ‘cortical FFRs (cFFRs)’ from here on. As is the case for scalp-recorded FFRs, the cFFRs closely followed the Mandarin tones’ fundamental frequency (Fig 1B). All four Mandarin tones elicited robust cFFRs (Fig 1C & 1F) in the electrode trajectories that were inserted along the HG, PT, and STG. The cFFRs that showed the highest amplitudes and signal-to-noise ratios were found in the electrode contacts closest to HG (Fig 1A & 1E) (*p*<0.05, Permutation based ANOVA followed by post-hoc paired t-tests on bootstrapped trials).

The magnitudes of the cFFRs were highest for tone stimuli with lower-F0, i.e., most robust in T3 (89-111 Hz) and T2 (109-133 Hz), followed by T1 (∼129 Hz) and T4 (140 to 92 Hz) (Fig 1D). This pattern is clearly visualized within the cFFRs to T2 and T4, where strong inter-trial phase coherence (ITPC) or phase-locking can only be observed when the F0 of the vowel is low and phase-locking declines when the F0 is high (Fig 1D & G).

### Cortical FFR latencies in human sEEG do not reflect volume-conducted activity from the brainstem

We assessed the latencies of the cFFRs for vowel T3 as they were the strongest and present throughout the stimulus duration. The latencies of the onset portion of the cFFRs were 14-16 ms, which is much later than the latencies expected of brainstem responses. Similar to the onset latencies, cFFR latencies (based on cross-correlation lags with maximum correlation coefficient) in the right HG (Liégeois-Chauvel et al., 1994) were ∼13-26 ms. These latencies are not consistent with the earlier neural conduction delays expected of inferior colliculus activity and suggest that the recorded cFFRs reflect phase-locking of post-synaptic potentials in cortical neurons.

### Hemispheric asymmetry in cFFRs

Hemispheric asymmetry was analyzed in Hum1 with bilateral temporal lobe coverage. The high-quality and high signal-to-noise ratio FFR data allowed us to statistically assess hemispheric asymmetry within the subject. cFFRs to the Mandarin tones showed a distinct hemispheric asymmetry, consistent with a prior study using MEG (Coffey et al., 2016). The electrodes in the right hemisphere showed higher amplitude cFFRs to the Mandarin tones (*p*<0.01, permutation-based t-tests on signal-to-noise ratios on bootstrapped cFFR samples). The rightward symmetry was also seen in the ITPC spectrograms and pitch tracking accuracy to the Mandarin tones (Fig 1D & 2B), which together indicate better phase-locking to the stimulus F0 in the right hemisphere. The better phase-locking in the right hemisphere was also seen in an additional set of non-speech stimuli i.e., click trains with repetition rates in the human pitch range (Supplementary Fig 1). We used a Hidden Markov Model (HMM) to decode the mandarin tones from the cFFRs. The cFFRs from the right hemisphere tracked the stimulus pitch better than in the left hemisphere (Fig 2b). Consequently, decoding accuracies were higher in the right hemisphere than the left hemisphere (Fig 2C). The pattern of tone decoding errors (‘confusions’) correlated significantly (p<0.05) between cFFRs from the right hemisphere and the scalp FFRs from a set of 20 subjects, but the same was not true for the cFFRs from the left hemisphere and the scalp FFRs (Fig 2D), Despite this difference, multidimensional scaling analysis revealed similar clustering of tone FFRs across the scalp, right HG, and left HG (Fig 2E).

**Figure 2.**
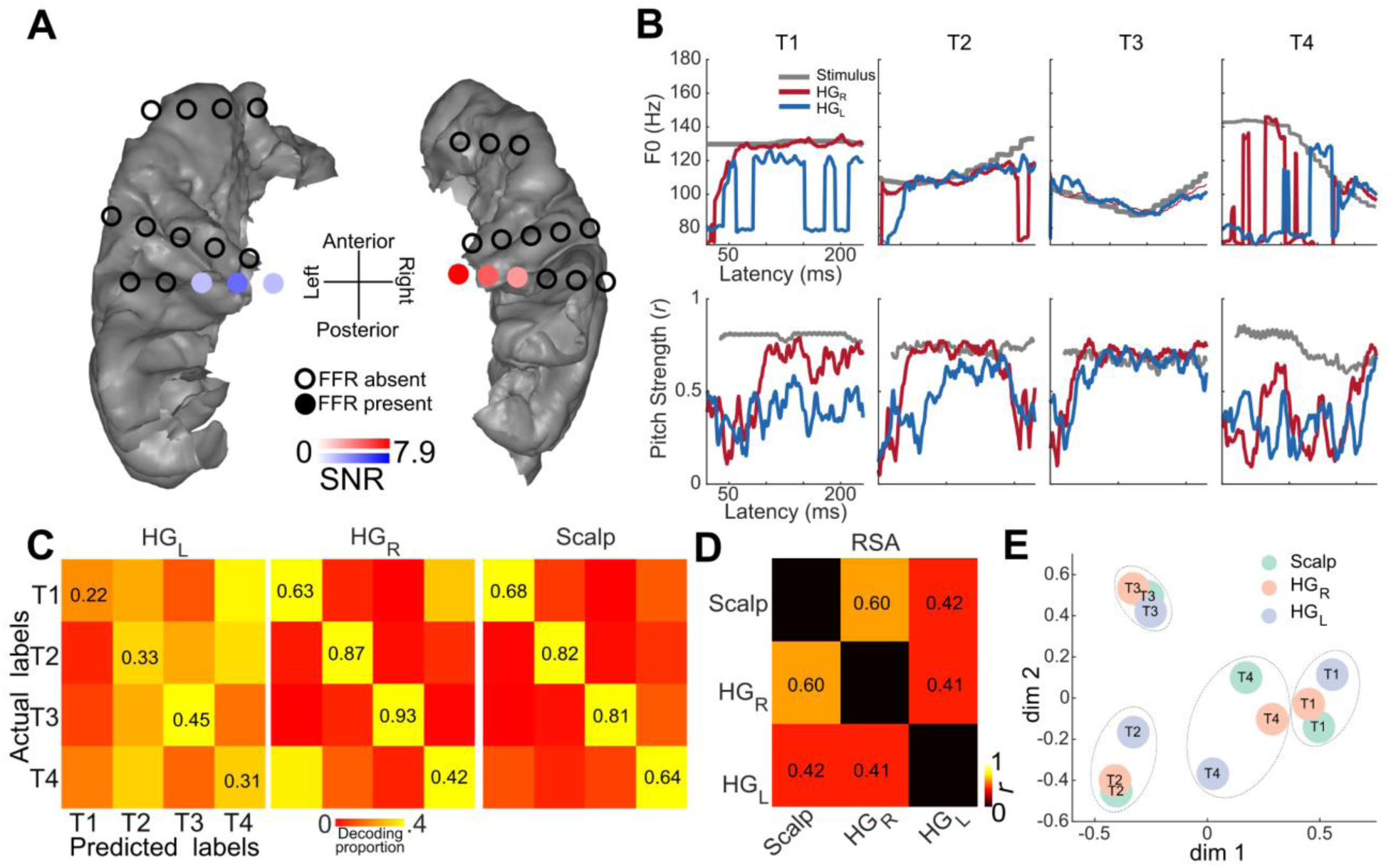
Hemispheric differences in the frequency following responses (FFRs) in Hum1. **A.** Electrodes implanted in the superior temporal plane are shown with filled circles marking significant FFR root mean square amplitudes above the pre-stimulus baseline (p<0.05 permutation-based bootstrapped t-tests) and unfilled circles mark electrodes where the FFRs were not significantly above the baseline. **B.** Pitch tracking measures (sliding window autocorrelation analysis) of the FFRs from the best electrodes in the two hemispheres showing the hemispheric differences. Top row shows the derived F0 track in Hz (best lag in each sliding window) in the right (HG_R_) and left (HG_L_) heschl’s gyri. Bottom row shows the pitch strength (maximum correlation coefficient in each sliding window). **C.** Hidden markov model based decoding accuracies of the Mandarin tone from the FFRs recorded in the right and left Heschl’s gyri from the sEEG recording, scalp EEG recordings from 20 different participants. **D.** Representational similarity (Pearson’s correlation) between the FFR confusion matrices (diagonals removed) in C. The correlation between HG_R_ and scalp FFR was significant (p=0.039) while the other correlations were not significant. **E.** Multi-dimensional scaling of the confusion matrices in C, visualizing the confusion space for decoding the Mandarin tone.

### Representational similarity analysis (RSA) of cross-species and cross-level FFRs

As in the humans, we recorded EEG in both animal model systems in order to establish scalp-derived FFRs as a translational bridge between the three species. Recordings in the animal models used the same Mandarin tone stimuli previously used for the human subjects. The scalp-recorded FFRs in both model species showed FFR activity (Fig 1I, 1L, & 1N) above the pre-stimulus baseline (SNR_Maq_ = 3.2, SNR_GP1_ = 7, SNR_GP2_ = 3.4) and showed the expected phase-locking to the F0 of the stimuli (Fig 1J, 1M, & 1N). Both species showed FFRs that correlated (*r_Maq_* = 0.45, *r_GP_* = 0.48, *r_GP1_* = 0.5, *r_GP2_* = 0.5, *r*-maximum cross-correlation coefficient) with the stimulus at latencies (Lat*_Maq_* = 3ms, Lat*_GP_* = 3.5ms) expected of early brainstem responses.

Furthermore, we recorded local field potentials (LFPs) from electrodes in PAC to compare against the intracranial LFP recordings in the human epilepsy patient. Similar to the scalp-recorded FFRs, the intracranial LFPs in both model species also yielded strong amplitude cFFRs above the pre-stimulus baseline (SNR_Maq_ = 18.3, SNR_GP1_ = 3.7381, SNR_GP2_ = 7.7) and readily showed the expected phase-locking to the F0 of the stimuli (Fig 1J, 1M, & 1N). The latencies of the cFFRs (Lat*_Maq_* = 11.6ms, Lat*_GP1_* = 9.7ms, Lat*_GP2_* = 10.1ms), however, were longer than scalp-recorded FFRs in both species (*p*s<0.001 in both GPs and Maqs, on sign rank comparison of stimulus to response cross-correlation latencies on bootstrapped samples).

We used RSA to quantify similarities between humans and animal models across different recording levels (intracortical vs. scalp) (Fig 3). RSA was performed on confusion matrices constructed from FFRs recorded using harmonized stimuli (four Mandarin tones) across-species and levels. Human scalp data were derived from FFRs recorded in 20 participants from a previously published study (Reetzke et al., 2018). In the macaque and GP subjects, scalp FFRs were recorded from cranial EEG electrodes surgically implanted in the skull. In the macaque, intracranial data were recorded from an electrode positioned immediately above layer 1 of the primary auditory cortex (PAC) from a chamber implanted over the frontal cortex. In the GP, intracranial data were recorded from an electrode contact estimated to be positioned in putative layer 4 of PAC. Visual inspection shows that the pattern of phase-locking of FFR and cFFR in the animal models was similar to that seen in the human, with phase-locking declining rapidly with increasing stimulus F0 (Fig 1D, 1G, 1J, 1M, & 1N).

**Figure 3.**
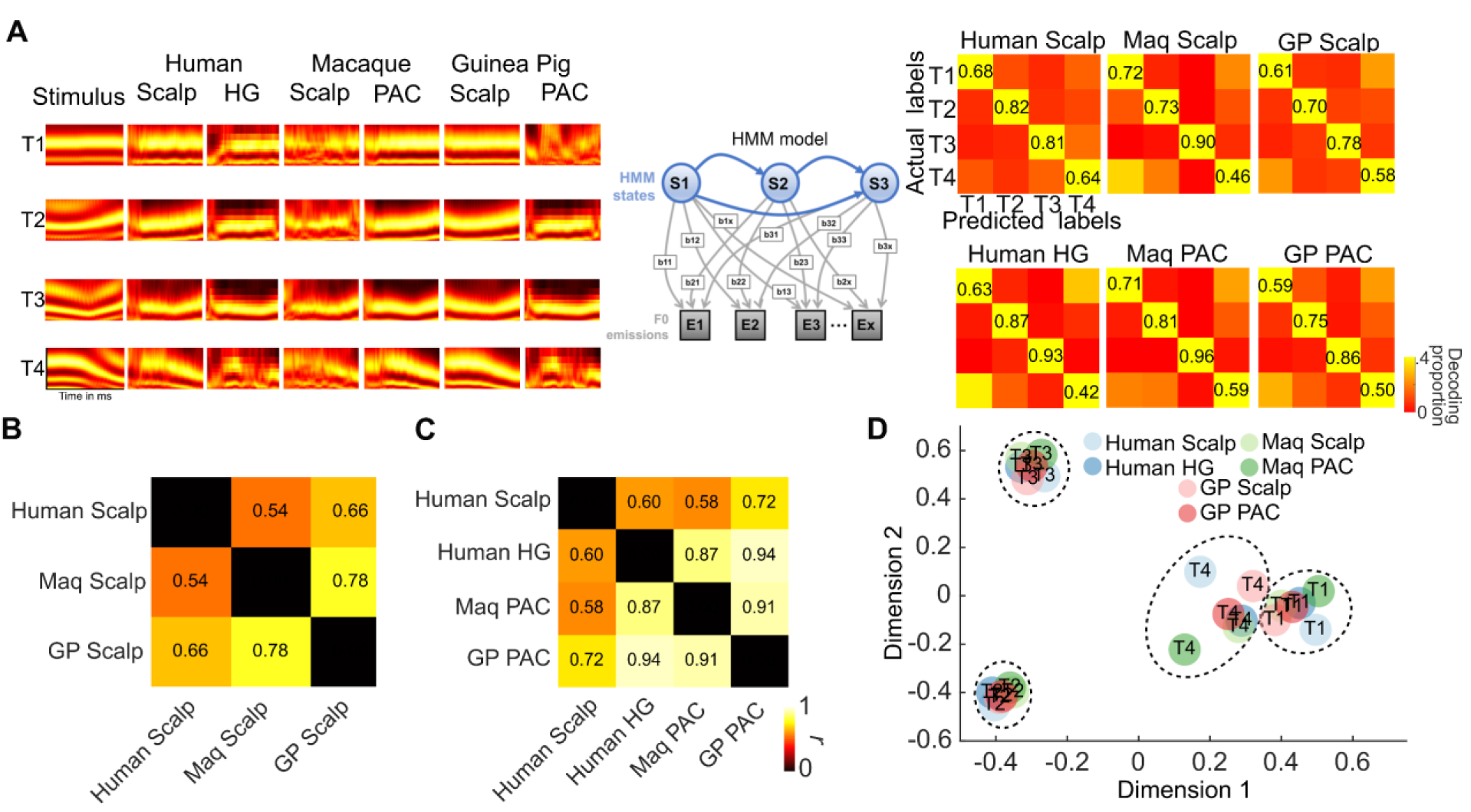
Cross-species and cross-level representational similarity analyses of FFRs. **A** Hidden Markov Model (HMM) was used to assess the extent to which pitch patterns (T1-High flat, T2-Low rising, T3-Low dipping, and T4-High falling) could be decoded from the FFRs. HMM decoding accuracies were estimated for each species (human, macaque, and guinea pig) and each level (scalp and intracortical). HMM decoding accuracies were significantly above chance for all species, levels. Confusion matrices *(right)* show the accuracies for decoding (along the diagonal) the pitch patterns from FFRs and error patterns. The averaging size used for decoding was adjusted to obtain comparable classification accuracies at the scalp and the cortex. The confusion matrices in A (principal diagonal removed) were assessed for correlations (Pearson’s *r*) between species and between levels to estimate the cross-species and cross-level representational similarity. **B.** shows the correlation between the confusion matrices for scalp FFR across species. FFRs recorded at the scalp share strong similarities (*ps*<0.05). However, the correlation between human and macaque scalp-recorded FFR did not reach statistical significance (p=0.06). **C.** shows the correlation between the confusion matrices of intracortical FFRs across species. Intracortical FFRs showed very strong correlations (*ps*<0.05). Also shown is the correlation of intracortical FFRs across species and the scalp-recorded FFRs in humans. The human scalp FFRs and GP intracortical FFRs did not show a significant correlation (p = 0.049). . **D.** Multi-dimensional scaling (MDS) analysis of the pitch patterns based on the confusion matrices in each species and level.

We decoded the Mandarin tone categories from scalp and intracranially recorded FFRs for all species using an HMM classifier (Llanos et al., 2017; Reetzke et al., 2018). The HMM classifier performed at above chance levels (>0.25) across species and levels in identifying the correct pitch patterns from the FFRs (principal diagonal, Figure 3A). Human and animal FFR confusion matrices were strikingly similar at both the scalp (Fig 3B) and intracortical levels (Fig 3C) (p<0.05 on Pearson’s correlation of confusion matrices without the principal diagonal) with stronger similarity seen for the intracortical cFFRs (Fig 3C). However, the scalp FFRs also yielded subtle species-specific differences (with greater similarity between human and GPs, relative to the macaque model).

### High-density intracortical recordings in animal models reveal the laminar distribution of cFFRs

Although intracortical recordings from human subjects provide high spatial and temporal resolution, they are still prone to contamination by volume-conducted fields from the brainstem and subcortical nuclei, and do not provide cortical layer-specific information. To overcome this limitation, we turned to laminar recordings from multi-contact electrodes traversing all layers of PAC approximately perpendicular to the cortical sheet (Fig 4) in the two animal models. These recordings allowed us to compute current source densities (CSD), which reflect post-synaptic currents and the corresponding passive return currents in the local cortical populations. Current sinks and sources are independent of volume-conducted potentials from the brainstem and the midbrain. The CSDs can be used to determine whether the post-synaptic currents in cortical populations are phase-locked to the stimulus, and if so, at which cortical depth and latency do they arise. In addition, these laminar recordings also allowed us to assess the prevalence, latency, and cortical depth of multi-units that phase-lock the F0 of the stimulus.

**Figure 4.**
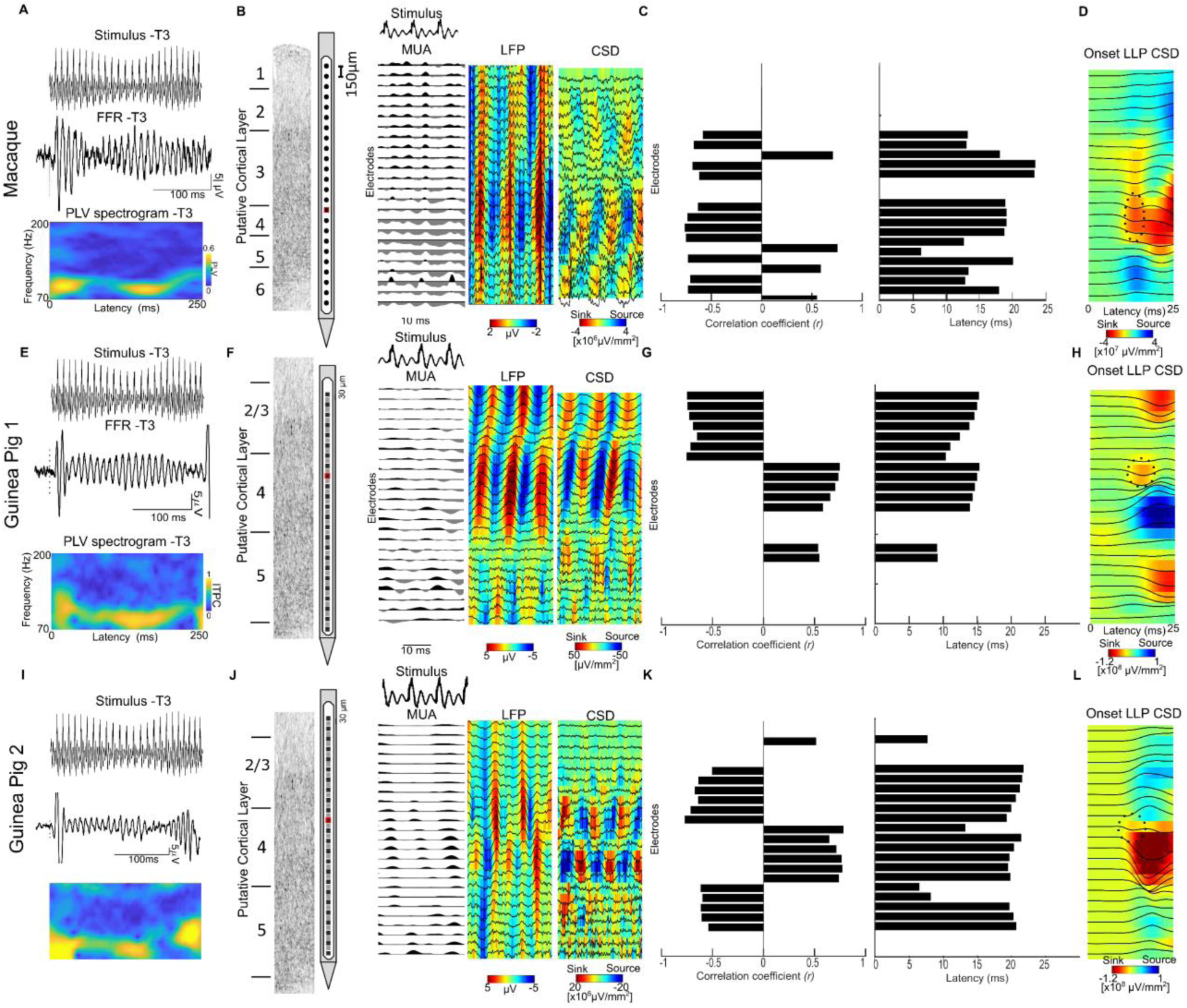
Cortical source for FFRs to speech syllables in the macaque and guinea Pig models: FFRs recorded in the macaque and guinea pig models using depth electrodes. **A**, **E** & **I** show sample stimulus (T3-low frequency dipping contour) and FFRs with the best signal to noise ratios in the putative layer IV in the laminar probe. Waveforms shown are from the electrode with maximum amplitude. Also shown is the phase-locking value spectrogram for the FFR (electrode shown in red in B and F). **B** & **F** show FFRs to stimulus T3, recorded using multi-channel laminar-depth electrodes in the macaque and the guinea pig, respectively. The FFRs are shown only for a short segment of the stimulus to clearly visualize the patterns. The multi-unit activity (MUA) and local field potentials (LFP) show strong phase-locked activity to each periodicity cycle in the stimulus. The source-sink configuration in the reference-free current source densities (CSD) show existence of an FFR generator in putative Layers III and IV of primary auditory cortex in all three animals. **C**, **G** & **K** show the stimulus to responses correlation coefficients and latencies (lags) between the stimulus and the FFR CSDs in each layer. The shift in polarity of the CSDs is manifested as a shift in the sign of correlation coefficient and shows sources of FFRs in two cortical layers. **D, G, &** L show the current source densities (CSD) of the long latency potentials (LLPs) to the onset of the Mandarin tones. The first sink (dotted ellipse) in the CSD marks the putative cortical layer IV.

Fig 4D, 4H and 4L show CSDs of the low-pass filtered local field potentials (1 – 70 Hz) aligned to the stimulus onset. This analysis identified expected patterns of sources and sinks for both animals that were used to identify the putative location of thalamorecipient cortical layers (layer IV and deep layer III), as well as supra- and infra-granular layers. We then computed the CSDs using the same filter setting used for the FFRs. Fig 4B & F show a 30 ms long snippet of the CSD FFRs from the sustained portion of tone 3 for both species (Maq2, GP1, and GP2). Note the presence of several currents that entrained to the F0 of the stimulus (stimulus to CSD correlation >0.5, Fig 4C, 4G, & 4K). We will refer to these currents as cortical frequency-following currents (cFFC). The most prominent cFFC was located in putative thalamo-recipient layers, and a second, somewhat weaker, cFFC with opposite polarity was identified in infra-granular layers. There was also an indication of a third and even weaker opposite polarity cFFC in supra-granular layers.

The cFFCs in both species showed a strong correlation with the stimulus at latencies of 12-25 ms in the macaque, and ∼10-25 ms in the GPs (Fig 4C, 4G, 4K). These latencies are consistent with a cortical origin. Only one infragranular cFFC in macaque had a latency of 6.3 ms that seemed inconsistent with a cortical origin. It is likely that this particular cFFC does not exclusively reflect post-synaptic activity, but rather a very large-amplitude spike that was isolated at this electrode contact and was bleeding into the frequency range of the FFR. Given the short latencies, it is likely that the spike in question corresponded to a passing thalamocortical fiber, rather than an infragranular cortical neuron.

In both species, the strongest and most prominent cFFCs were recorded in granular layers, and most likely reflect active postsynaptic currents in response to F0-locked thalamic input at basal dendrites (Fig 4B, 4F, 4J). It is less clear if the cFFCs in infra and supra-granular layers reflect active postsynaptic currents which might be indicative of the propagation of phase-locked responses to these layers or if they exclusively reflect passive return currents. In order for the phase-locked activity to spread beyond thalamo-recipient layers, not only the post-synaptic input currents but also the output, i.e., their firing rates, would have to be entrained to F0. We thus assessed the frequency following in the multi-unit activity (MUA), a measure of neural firing rate of small clusters of neurons in the immediate proximity of the electrode contact in the thalamo-recepient layers.). Electrodes with MUA showing stimulus to response cross-correlation coefficients>0.5 were present in both animal models (*r_Maq_* = 0.53, Lat_Maq_ = 13.5 ms, *r_GP1_* = ∼0.53, Lat_GP2_ = ∼13ms*, r_GP1_* = ∼0.6, Lat_GP2_ = ∼13ms). The presence of FFRs in the MUA suggest that the thalamo-recipient layers not only receive phase-locked input but also fire in a phase-locked manner to the stimulus F0 in both animal models. These results indicate that the thalamo-recipient layers not only receive strong phase-locked input from the thalamocortical fibers, but may also propagate the FFRs to downstream cortical layers, albeit with reduced phase-locking strength.

### Relationship between scalp and cortical FFRs

Because both intracranial and scalp FFR recordings were obtained in the same macaque and GPs, we used the opportunity to examine the power and latency across frequencies of the scalp and cortical FFRs to infer similarities. The cortical FFRs were higher in amplitude than the scalp FFRs, presumably due to the proximity of the electrodes to the cortical sources. The comparison of the spectral characteristics of the scalp and cortical FFRs were thus made by normalizing the spectral estimates. Compared to the scalp FFRs, the cFFRs from the PAC were predominantly composed of low-frequency F0 energy relative to higher harmonics (Fig 1 and 5B). Fig 5A shows the FFRs to tone 3 (low-frequency dipping contour) recorded from the scalp and cortex in the macaque and the GPs. Cortical cFFRs in both species showed longer latencies than the scalp-recorded FFRs (*p*s<0.01 permutation based Wilcoxon sign rank tests on bootstrapped FFR trials) (Fig 5A). While the phase-locking of cFFRs to Mandarin tones were higher in the PAC than at the scalp (Fig 1), the decline in phase-locking with increasing frequency was similar at both the PAC and the scalp. This can also be seen in the difference in normalized power spectral density between the scalp and cortical FFRs at the high frequencies when normalized based on maximum spectral amplitude (Fig 5B).

**Figure 5.**
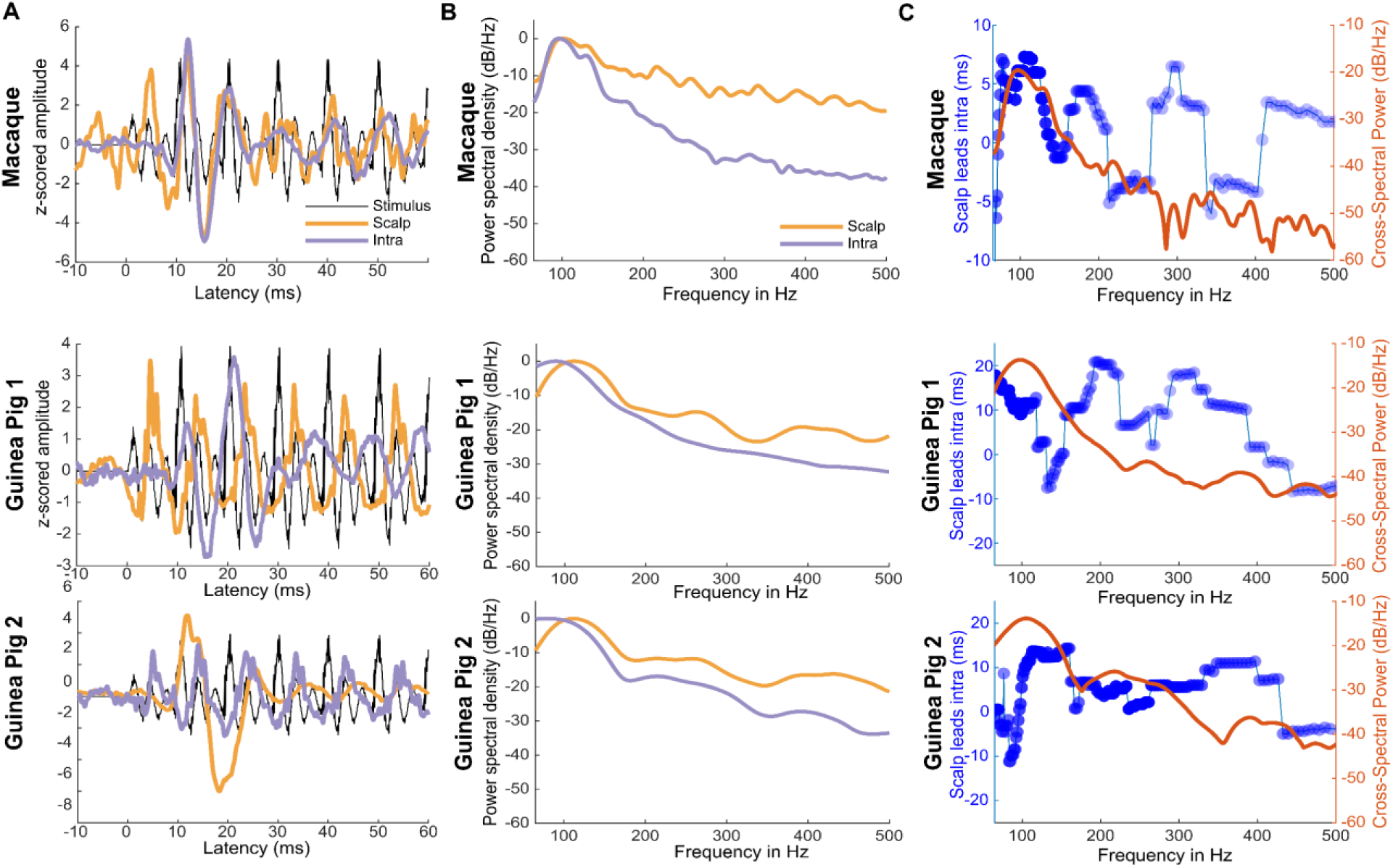
Scalp and intracortical FFRs in the macaque and the guinea pig. **A.** FFR waveforms to Mandarin tone 3 (low dipping F0) at scalp and the cortex in the macaque and the two guinea pigs. **B.** Normalized power spectral density of the scalp and intracortical FFRs in the macaque and guinea pig, showing the different the low-frequency dominance of the intracortical FFRs. **C.** Difference between scalp-recorded and intracortical FFRs. Blue tracings show the latency between scalp and intra-cortically recorded FFRs. Circles with darker colors (colors normalized to maximum correlation across frequencies)) indicate a higher correlation between the scalp and intra-cortically recorded FFRs. Intracortical FFRs showed longer latencies than scalp-recorded FFRs at frequencies below 120 Hz. Red tracings show the cross-spectral density between scalp and intra-cortically recorded FFRs showing shared power at low frequencies.

Cross-spectral power analysis revealed that scalp and cortical FFRs shared strong power coherence with the stimulus near the F0 (70-110 Hz), which declined rapidly at higher frequencies in the cortical FFRs relative to the scalp FFRs (Fig 5C). This trend was similar in all animals (Maq1, GP1, and GP2). This pattern indicated that the FFRs recorded at the scalp and the cortex were similar in power spectral density at the low-frequency regions. Such a pattern can be caused either by a single common source or by more than one source with similar spectral properties but different temporal properties. We thus estimated the cross-correlation strength and latency between the scalp and cortical FFRs across frequencies in the time-frequency domain. The maximum correlation between the scalp and cortical FFRs was seen at frequencies <120 Hz. However, at these frequencies, the cortical FFRs showed delays of∼7 ms in the macaque, and ∼10 ms (Fig 5C) in the GP in comparison to the scalp FFRs. This indicates the presence of temporally disparate but spectrally overlapping neural sources of the scalp-recorded FFRs. These delays can also be prominently seen in the latencies of the cFFRs, which are higher than conventionally obtained scalp-recorded FFRs (Fig 5C)

We further applied a blind source separation approach to disentangle the spectro-temporally overlapping components that contribute to the scalp-recorded FFR. We used independent component analysis (ICA) as the source separation approach (Maq - Fig 6 and GP- Fig 7). In the macaque, we submitted all the scalp (33 electrodes) and intracortical electrodes (96 electrodes spanning the superior temporal plane, prefrontal, and pre-motor cortex to ICA decomposition. We extracted 12 ICAs that explained 96% of the variance. Among these, we focused on four ICAs each of which explained >10% of the variance individually, and who as a group explained 75% of the variance (Fig 6). ICA2 was consistent with a volume-conducted generator from the regions distant to all intracranial electrodes (Fig 6A). This is apparent from the widely distributed ICA weights across electrodes. Furthermore, ICA2 had a latency of 3.4 ms and showed prominent power coherence with the stimulus at F0 as well as the higher harmonics (Fig 6C). In contrast, ICA1 had a longer latency (18.1 ms), responded strongest to the F0, and exhibited a steeper gradient between electrodes in the superior temporal plane and motor/premotor cortex. The power coherence with the stimulus also declined in frequency faster than in ICA1. These three findings are consistent with a generator in the primary auditory cortex. This putative cortical ICA1 also propagated to the scalp and contributed to the scalp-recorded FFRs. Similarly, topographies and latencies of ICA4 (lat=0.5 ms suggested a cortical origin and propagated to the scalp electrodes. ICA3 and ICA4 in contrast to ICA1 show spatial weights that are opposite in polarity and hence possibly emerged from different cortical sources with different orientations.

**Figure 6.**
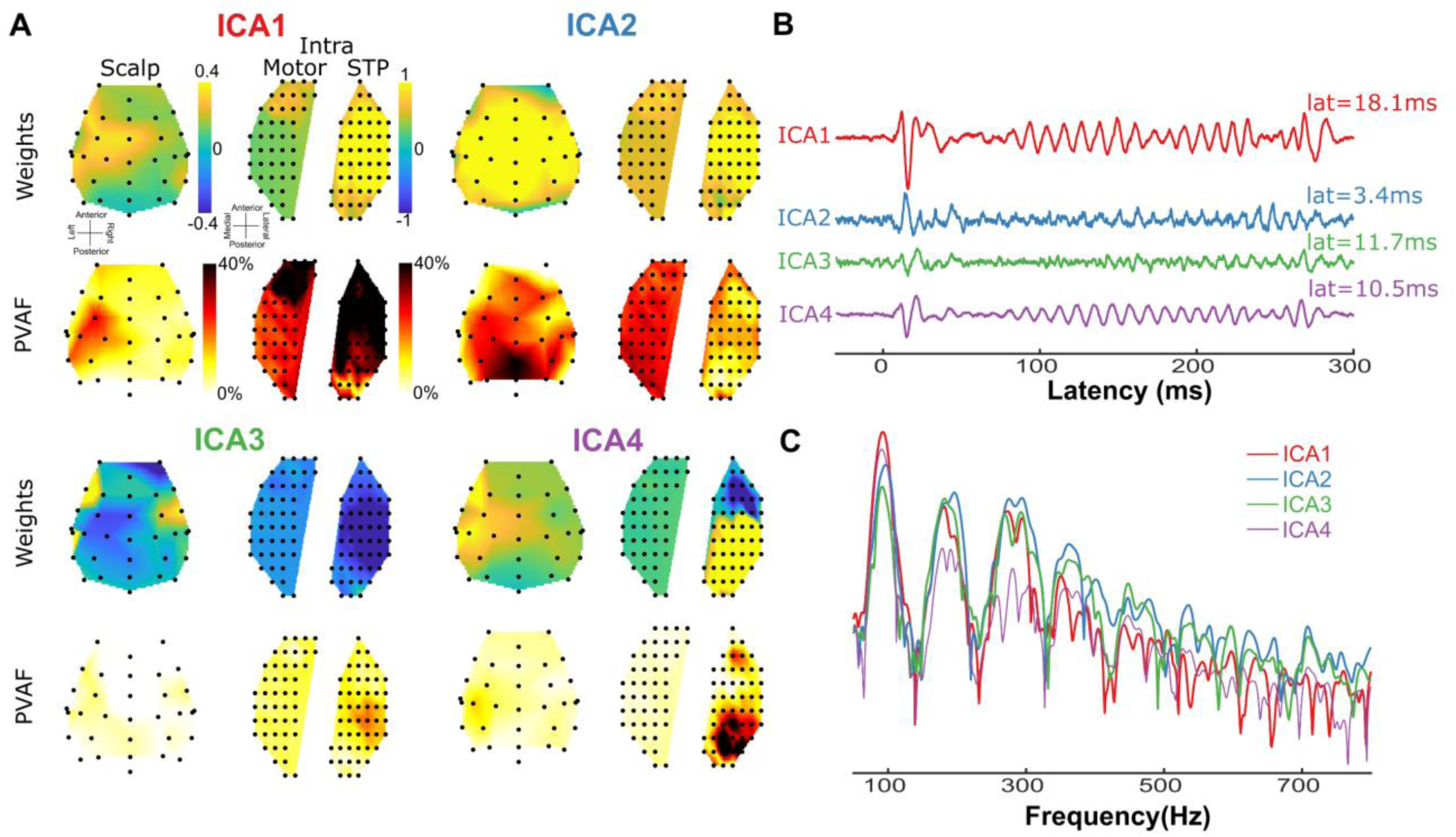
Evaluating the contribution of intracortical FFRs to scalp FFRs using independent component analysis (ICA) in the Macaque. **A.** Spatial loadings (ICA weights) of the top 4 independent components onto the scalp and intracortical (Intra electrodes located in the motor regions and in the superior temporal plane) electrodes and the percentage variance accounted for (PVAF) by the ICAs at each electrode. **B.** Time course of the activations of the top six ICs (amplitude in arbitrary units) for stimulus T3 (low dipping F0). **C.** Power spectral coherence of the top six ICAs contributing to the FFRs(amplitude in arbitrary units).

**Figure 7.**
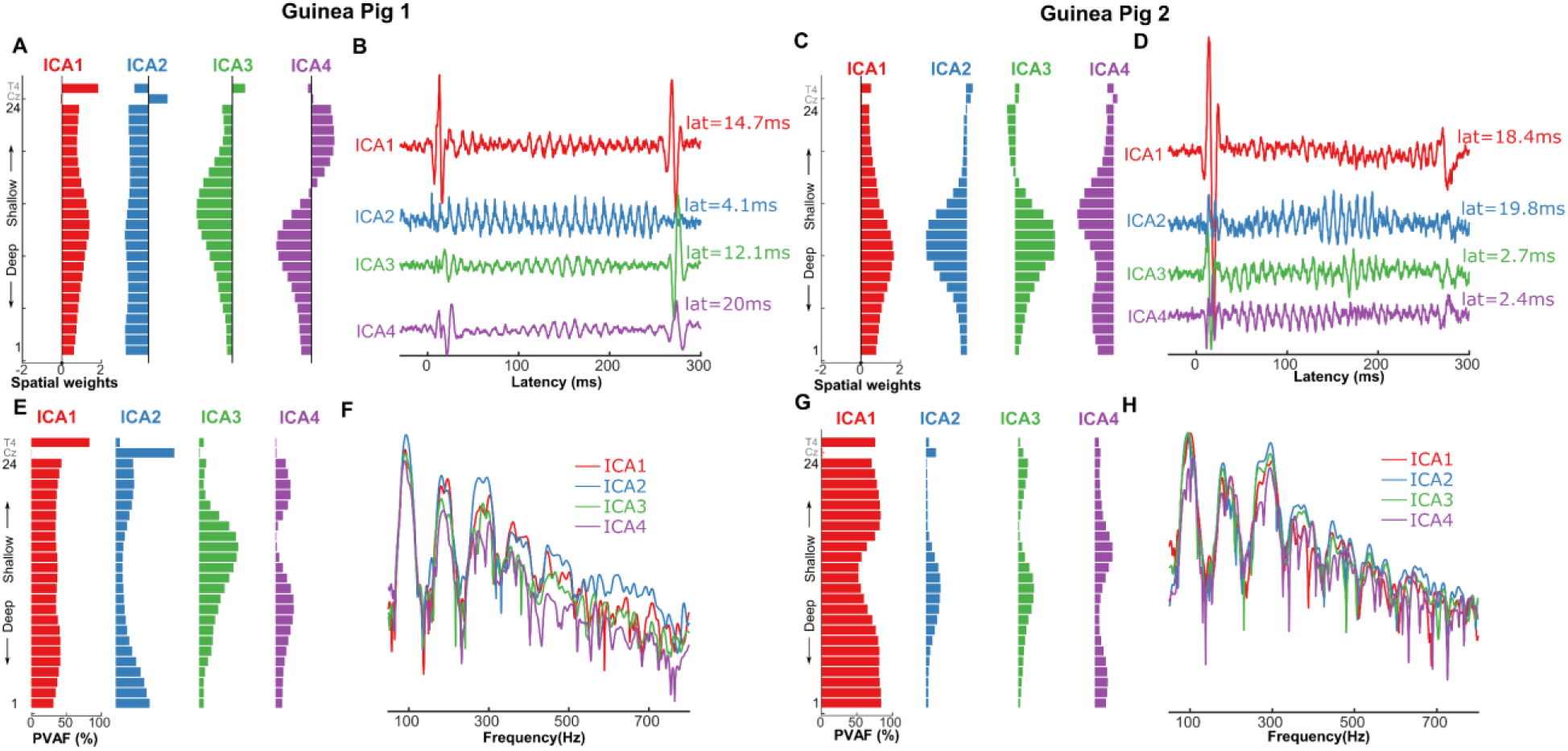
Evaluating the contribution of intracortical FFRs to scalp FFRs using independent component analysis (ICA) in the GP. **A.** Spatial loadings of the top 4 independent components onto the intracortical laminar electrodes (1-24) and scalp electrodes (Cz and T4). **B.** Time course of the activations of the top four ICs (amplitude in arbitrary units). **C.** Spatial loadings of the top six independent components onto the intracortical laminar electrodes (1-24) and scalp electrodes (Cz and T4). **D.** Power spectral coherence of the top four ICAs contributing to the FFRs (amplitude in arbitrary units).

In the GP, we submitted the averaged FFRs for each of the Mandarin tones at the 24 electrodes along the layers of the PAC, and 2 electrodes placed on the scalp to ICA. The scalp electrodes were placed on the vertex of the scalp (Cz : midpoint of the head along both sagittal and coronal axes) and on the temporal surface of the scalp close to the auditory cortex (T4). Six ICAs were extracted that explained ∼99% of the variance in the FFRs. Among these, the first four components explained >10% of the variance in the FFR amounting to a total of 93.5% (GP1) and 99% (GP2) (Fig 6). Based on visual inspection, the weights of ICA2 in GP1 were not modulated appreciably along the laminar electrode layout. Similarly, ICA3 and ICA4 in GP2 were not modulated appreciably along the laminar electrode layout. This is consistent with volume-conducted activity from distant brain regions, in this case most likely the brainstem (Fig 7A & C). It should be noted that the spatial loadings of the volume conducted ICs in GP2 largely follow the same trend as GP1 but are not exactly similar. This could be in part influenced by the large cortical onset and offset responses that propagate to the scalp in GP2 which are not as apparent in GP1.

In GP1 the putative subcortical ICA2 had a latency of 4.1 ms and showed strong power coherence with the stimulus F0 as well as the harmonics (Fig 7F). This putative subcortical ICA2 contributed almost the entire variance in the scalp electrodes placed at Cz, thus suggesting negligent contribution from other sources such as cortex. ICA1 (lat=9.7ms) and ICA3 (lat=12.3ms) showed maximum spatial weights around putative cortical layers 4 and 2/3 and contributed maximally to the variance at the electrode T4 that was placed very close to the surface of primary auditory cortex. In contrast, it did not contribute to variance at the scalp electrode Cz. While ICA4 (lat =15.1ms) was also consistent with a cortical source, it contributed to very little variance in the scalp electrodes. Taken together, these results show that in the GP, the scalp electrodes placed at the midline are dominated by the subcortical sources, while those placed on the temporal scalp locations are dominated by cortical sources.

Similar results were also obtained in GP2 where the putative subcortical ICA3 and ICA4 had a latencies of 2.7 and 2.4 ms respectively, and showed strong power coherence with the stimulus F0 as well as the harmonics (Fig 7D). Unlike the volume conducted ICAs in GP1, the volume conducted ICAs in GP2 did not explain the entire variance in Cz. This could be a result of very large onset responses which skew the ICA decomposition. ICA1 (lat=18.4 ms) showed maximum spatial weights around putative cortical layers 4 and 2/3 and contributed maximally to the variance at the electrode T4 that was placed very close to the cortical surface of primary auditory cortex. In contrast, it did not contribute to the variance at scalp electrode Cz. While ICA2 (lat =19.8ms) was also consistent with a cortical source and contributed to the variance at the scalp, it consisted of a late onset portion and a cortical FFR portion that projects to both scalp electrodes. Taken together, these results show that in the GP animal model, the scalp electrodes placed at the midline are dominated by the subcortical sources, while those placed on the temporal scalp locations are dominated by cortical sources. It should be noted that the SNR of scalp electrodes in GP2 was lower than in GP1 due to the lower number of stimulus sweeps. Also scalp and intracortical FFRs in GP1 were recorded in separate sessions in GP1 while they were recorded simultaneously in GP2. These differences could have driven the subtle differences in ICA patterns between the two GPs.

Thus, the ICA results of both macaque and GPs suggest that the scalp-recorded FFRs contain weighted mixtures of both cortical and subcortical sources. The contributions of both these sources are dependent on the orientation of the net electrical dipole and the location of the scalp electrode. Regardless, the cortical contribution to the FFRs can be seen in both species, these cortical sources indeed emerge in the auditory cortex and propagate to the scalp.

## DISCUSSION

Research spanning three decades has richly characterized the properties of subcortical FFRs (Chandrasekaran and Kraus, 2010a; Skoe and Kraus, 2010a; Krizman and Kraus, 2019; Coffey et al., 2021), but open questions remain with respect to characteristics of the cortical FFR sources and their contribution to the scalp recorded FFRs. Our study establishes a cross-species (human, macaque, GP) and cross-level (intra-cranial, scalp) platform to study cortical FFRs and their contribution to the scalp-recorded FFRs. We present several novel results; (1) All species readily exhibited FFR-like responses to Mandarin stimuli in both scalp and cortical recordings. (2) Better encoding of lower frequencies and longer latencies was a characteristic of the cortical FFRs in all species. (3) The bilateral FFR recordings from the Heschl’s gyrus in humans showed robust encoding and representation of higher fundamental frequencies in the right relative to left HG. (4) RSA revealed striking similarities in the cortical representation of F0 contours, firmly establishing the macaque and guinea pig as viable animal models to study the cortical FFRs to human speech. (e) Laminar recordings from the macaque GP auditory cortices demonstrated the existence of cortical frequency-following currents (cFFC) to human speech sounds in thalamo-recipient layers of PAC. (f) Using EEG and large-scale intracranial recordings in the same animal model we traced the putative contribution of the auditory cortex to the scalp-recorded FFR. Taken together, our results provide novel insights into the properties of the cortical source of the FFRs to time-varying pitch contours. In the sections below, we highlight and expand upon the key findings within and across species.

### Cortical FFRs in humans show distinct low-frequency and right hemisphere bias

Previous studies used distributional source modeling applied to EEG or MEG data to study the cortical source of FFRs (Coffey et al., 2016; Bidelman, 2018; Hartmann and Weisz, 2019). Here we circumvented the challenges of inverse source localization by using direct intracranial recordings in two human participants, and confirmed that robust cortical FFRs to pitch patterns could be evoked in the Heschl’ gyri. These cortical FFRs phase-locked only to the stimulus fundamental frequency, while subcortical FFRs can track speech harmonics as high as 950 Hz (Galbraith et al., 2000; Plyler and Ananthanarayan, 2001). Further, the latencies of cortical FFRs (17-23 ms) were significantly longer than expected of subcortical FFRs (Du et al., 2009; Wang and Li, 2018). Compared to earlier studies, we examined the cortical FFRs to higher F0s (made possible by intracranial recordings), and showed cortical FFRs to F0s as high as 150 Hz.

Bilaterally, the electrodes in the HG showed substantially stronger FFRs compared to those in the PT. No other cortical regions close to the HG showed FFRs. The PT did not phase-lock to the periodicity of the stimulus, which might indicate a transformation of temporal pitch code into a place or a rate-place code in the auditory-association cortex. This pattern was also consistent in the macaque data (Maq1) where only electrodes closest to the primary auditory cortex showed strong FFRs. Weaker FFRs on electrodes in motor, pre-motor and prefrontal cortex were likely volume-conducted fields not originating in the motor regions.

Consistent with previously reported rightward bias in the cortical FFR activity (Coffey et al., 2016, 2017a, 2017b; Gorina-Careta et al., 2021), we found evidence of distinct rightward asymmetry of FFR magnitudes and steeper phase-locking decline with F0 in the left compared to the right HG. The right hemisphere asymmetry observed in our study may underlie processing differences of melodic and prosodic features in (non-native) speech and music (Zatorre and Belin, 2001; Coffey et al., 2017b).

Experience-dependent changes in FFRs to Mandarin tone stimuli have been extensively used to inform theoretical models of subcortical plasticity (Patel and Iversen, 2007; Krishnan et al., 2012; Skoe and Chandrasekaran, 2014). Our finding of a cortical contribution to FFRs elicited by these very same stimuli adds important new information directly relevant to these theoretical models. As a result, subcortical plasticity models based on the FFRs need to be revisited with a new lens that focuses on the relative cortical and sub-cortical contributions to experience-dependent plasticity. Given that macaque monkeys can be trained on various auditory tasks and given the similarities of human and monkey FFRs, they are a promising model species to quantify the relative cortical and subcortical contributions to emergent plasticity measured by the FFRs.

It should be noted that earlier intracranial human studies have shown the existence of FFRs to speech at the auditory cortex (Behroozmand et al., 2016; Guo et al., 2020). However, due to the coarse spatial resolution offered by human sEEG it cannot confirm the presence of cortical frequency following currents as against the thalamocortical input currents, which is essential to firmly establish cortex as a putative generator of scalp-recorded FFRs. By establishing similarities in FFR representation between the human and animal models, and by leveraging high-density laminar recordings in animal models, we were able to explore the laminar sources of the FFRs and breakdown the cortical contributions to the scalp FFRs with high spatial and temporal detail.

We leveraged representational similarity analyses (RSA) as a translational bridge across levels and species. This allowed us to further deep dive into the FFR sources in animal models at an fine anatomical resolution. Critically, despite differences in recording procedures, anatomy, and arousal states, we demonstrate strong similarities in representational structure between the cortical and scalp FFRs in both human and animal models. Further, the similarity across the species suggests similar representation of the F0 feature in the three species. Further, across the species, the falling tone (T4) was represented less robustly (more confusion) than the rising tone (T2) (less confusion) suggesting a cross-species similarity in preferential processing of stimuli with rising, relative to falling pitch (Peng et al., 2018). Due to these similarities, macaques and GPs may be well-suited to help answer important questions about the cortical FFRs. The extent of representational similarity across species was lower for scalp-recorded FFRs than intra-cortically recorded FFRs, which likely reflects a variability in dipole orientations of cortical FFR sources.

These results complement and expand an earlier study (Ayala et al., 2017), which explored the similarity of human and monkey scalp FFRs based on morphological characteristics of FFR to a single 40 ms /da/ syllable with a relatively steady F0. Going beyond a morphological comparison, we use a range of complex speech sounds with time-varying pitch to assess the species-specific similarity using RSA. We also establish homologies across three animal species along the evolutionary hierarchy, each of which can be leveraged to understand FFRs using advanced approaches that are species-specific; for example, optogenetic approaches can be efficiently used in guinea pigs to understand the effects of corticocollicular projections on FFRs, and macaques can be efficiently trained to categorize novel stimuli to reveal the effects of learning on FFRs. We have set a crucial template for future studies to examine the FFRs across species which is invaluable for leveraging species-specific analytical techniques to comprehensively understand FFR characteristics. Such comprehensive assays of the FFRs are vital to understand the factors underlying altered FFRs in various pathologies and to make the FFRs more readily interpretable and clinical viable.

A caveat regarding the RSA approach is that unlike the morphological analysis performed by Ayala and colleagues (Ayala et al., 2017) it is agnostic to subtle differences in absolute physical characteristics such as amplitudes, latencies and phases. This implies that the inference about similarity across species from the RSA alone should not be extrapolated to the specific physical characteristics of the FFRs.

### Cortical FFRs emerge in the thalamo-recipient layers of the primary auditory cortex

We consider FFRs as being of cortical origin if they arise from phase-locked post-synaptic currents in cortical neurons regardless of whether the post-synaptic currents are driven by thalamic or cortical input. Conversely, we define FFRs as having a subcortical origin if they arise from phase-locked post-synaptic currents of neurons located in subcortical nuclei. This working model is schematized in in Fig 8.

**Figure 8.**
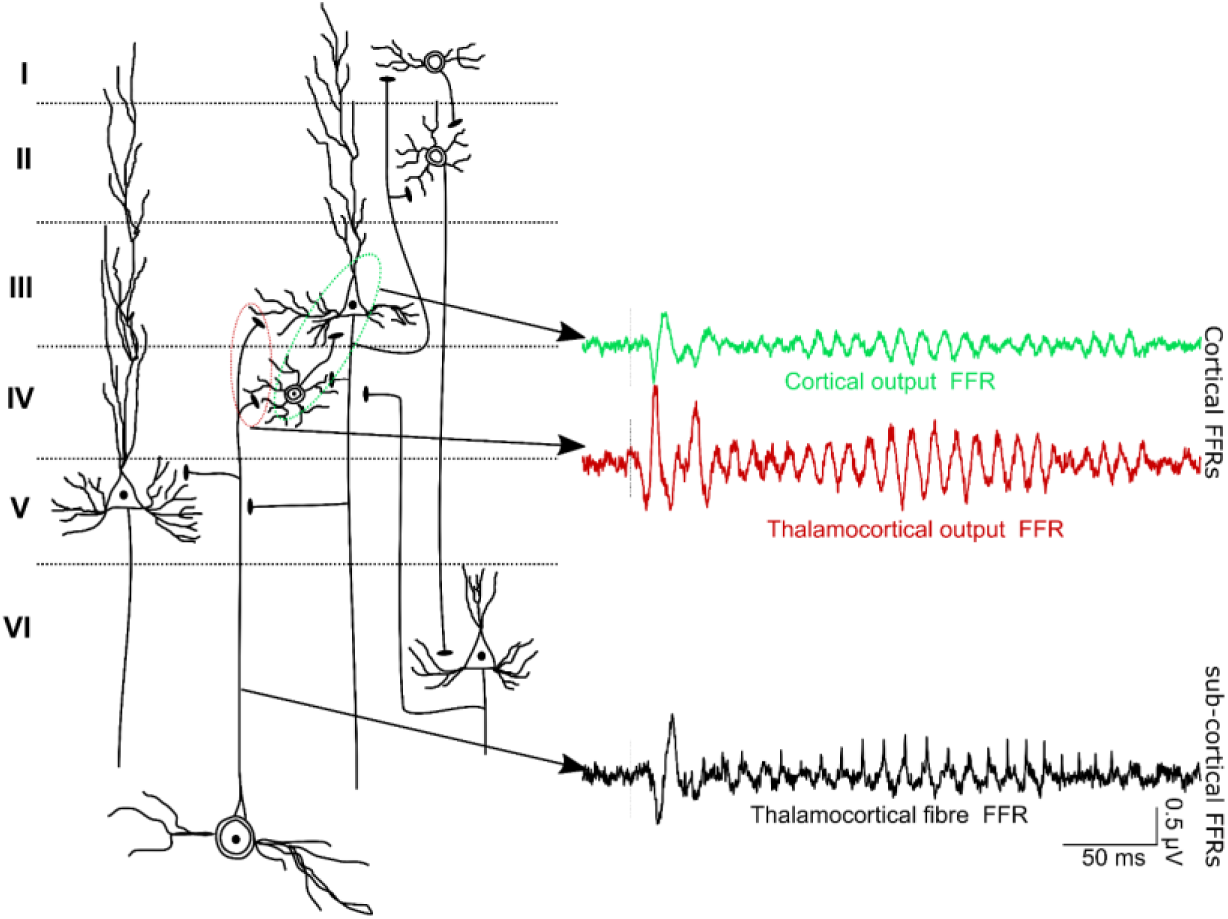
Schematic representation of the cortical FFRs and the subcortical FFRs. Any phase locked currents in the cortical layers are considered the cortical FFRs. Thalamocortical outputs that synapse with the layers III and IV in the auditory cortex generate post-synaptic potentials that are phase-locked to the stimulus F0.

With the laminar depth electrodes, we confirmed the existence of cortical FFRs across layers of the PAC. This was consistent in the macaque as well as both guinea pigs. We found that currents in thalamo-recipient layers follow the F0 input from the thalamocortical fibers. The outputs (MUA) of the thalamo-reciepient layers also follow the stimulus F0. Further, the FFR currents showed phase shifts across the different layers, suggestive of the electric field interferences that may dampen the net equivalent current dipole at the cortex. However, considering the lower strength of the cortical currents beyond the thalamo-recepient layer, it is parsimonious to assume that the thalamo-recipient layers primarily drive the FFRs recorded at the cortical surface in the human sEEG and macaque. While FFR currents have been demonstrated in earlier work (Steinschneider et al., 1999, 2003) for steady fundamental frequencies <100 Hz, here we show that cortical FFR currents show phase-locked activity as high as 150 Hz and follow the Mandarin pitch contours. These FFR currents could have partially contributed to the cortical FFR components found in earlier studies using non-invasive (Coffey et al., 2016, 2021; Bidelman, 2018; Gorina-Careta et al., 2021) and invasive assays (Behroozmand et al., 2016; Guo et al., 2020). However, it should be noted that the non-laminar assays (both non-invasive and invasive) could very well pick up both presynaptic and post-synaptic currents that effectively constitute the cortical FFRs (Fig 8). Nevertheless, we emphasize that the finding of cortical source of the FFR is not at odds with the well-established existence of subcortical sources in scalp-recorded FFRs (Marsh et al., 1975; Smith et al., 1978), but show that cortical sources also contribute to scalp FFRs

### Cortical contributions to extracranially recorded FFRs

The use of animal models enabled us to explore the similarities between the scalp and cortical FFRs in the same subjects. The power coherence between the scalp and cortical FFRs further shows that the cortical sources do not strongly contribute to scalp FFRs at higher harmonics. At the F0, however, there was a strong correlation between the scalp and cortical FFRs. However, the latencies of these correlations suggested that the cortical FFRs had longer latencies than the scalp FFRs.

We used independent component (IC) analysis to disentangle the contribution of the spatio-temporally overlapping cortical and sub-cortical FFR components and their contribution to the scalp recorded FFRs. We found short latency volume-conducted ICs that presumably emerged from the subcortical regions and propagated uniformly to all scalp and cortical electrodes. We also found strong ICs that were localized to thalamocortical recipient layers and projected to the scalp. These ICs likely reflect the bulk signal from the cortex that propagates to the scalp. However, not all putative cortical ICs contributed to the scalp-recorded FFRs due to dipole orientations that did not favor volume conduction to the scalp, and differed based on the electrode location. There was a very specific pattern of differential cortical contribution based on scalp electrode location. In the GP, the midline electrode predominantly picked up the subcortical component, while the electrode above auditory cortex predominantly picked up the cortical component. This effect is driven by the fact that in the GP the auditory cortex is largely on the surface of the temporal lobe with cortical layers oriented medio-laterally unlike humans and other primates where the auditory cortex is buried in the lateral sulcus with cortical layers oriented dorso-ventrally (Wallace et al., 2000; Petkov et al., 2006). This further implies that the contribution of the cortical FFRs, though smaller than the subcortical contribution, is present and dependent on the electrode montage used, the location of the auditory cortex and the folding patterns of the cortical surface. The use of two animal models and the multiple modes of intracortical recordings provided complimentary converging evidence for cortical source of FFRs. At the same time, it also sheds light on the species-specific differences and similarities that contribute better understanding of the FFR properties across animal models.

### Limitations and Future Directions

The sample size can be considered a limitation of the study, potentially limiting the generalizability of the study. However, it is very rare to record FFRs from bilateral Heschl’s gyrus in the same human subject, and we statistically show the comparison between the two hemispheres at a single subject level. Similarly, we show high quality replicable recordings in the two macaques and the two guinea pigs. Further, we show that the finding of FFRs to speech is localized to the HG similar in both our human subjects. In addition, the results of both GPs are also largely similar. We also report results of individual human, macaques, and guinea pigs for better transparency and to avoid over-generalization across the small sample sizes. We also offer several complementary analyses within and across species to facilitate our analysis which show converging evidence for our interpretations. As in all intracranial explorations in human participants, the results are still based on human brains with atypical brains with epileptic seizures, which can hinder generalizability to typical human participants. The macaque and guinea pig models provide important translation bridge in understanding the laminar profile and the cortical contribution to the scalp FFRs. However, the differences in cortical folding properties and anatomical orientation of the brainstem and the auditory cortices could potentially confound generalization to human participants. We partially circumvent this problem by establishing homologies across the species for both scalp and intracranial FFRs. We did not have simultaneously recorded scalp FFRs in our human sEEG subjects due to clinical challenges, and leveraged scalp FFRs from a separate set of participants to understand the differences in scalp and cortical FFRs. Future studies using simultaneously acquired scalp and cortical FFRs will be invaluable in directly inferring about the cortical contribution in human subjects. Further extensive and systematic characterization of the scalp and cortical FFRs would help in better translation of cognitive, and behavioral interventions on the neural plasticity at the cortical vs subcortical levels.

## CONCLUSIONS

We demonstrate the existence of a cortical source of FFRs using direct intracranial recordings to time-varying pitch patterns. Using an integrated cross-species approach with the macaque and GP models, we show laminar profiles consistent with a cortical source of the FFRs that emerge from the thalamo-cortical-recipient layers and not from the volume-conducted activity of subcortical neurons. We also show that while subcortical sources dominate the scalp-recorded FFRs, cortical sources do contribute to the scalp-recorded FFRs. Our research paves the way for a wide array of studies to investigate this cortical FFR source’s relevance in auditory perception and plasticity.

## ACKNOWLEDGEMENTS

This investigation was supported by National Institute of Health funding to R01-DC013315 to B.C., S.S., T.A, and T.T; RF1-MH114223 to T.T., and R01-DC017141 to S.S. This work was also supported in part by the resources provided through the University of Pittsburgh Center for Research Computing. Preliminary findings from this investigation were presented in the symposium of Advances and Perspectives in Auditory Neuroscience 2020.

## Supplementary Information Discussion: Laminar sources of FFRs

FFR current sinks were also observed in the deep layer V. It is not clear if these currents are just passive return currents or active currents in which case they would indicate the propagation of phase-locked activity to layer V neurons which give rise to cortico-collicular feedback projections(Bajo and Moore, 2005; Bajo et al., 2010). Paradigms using stimulus probability effects have shown systematic effects on the FFRs, and have been considered as a putative marker cortico-collicular processing(Chandrasekaran et al., 2009; Skoe and Kraus, 2010b; Slabu et al., 2012; Gnanateja et al., 2013; Maruthy et al., 2017). The use of such paradigms in future studies using laminar recordings will help confirm whether layer V indeed underlies contextual-modulation of the FFR(Chandrasekaran et al., 2009; Skoe and Kraus, 2010b; Parbery-Clark et al., 2011; Slabu et al., 2012; Gnanateja et al., 2013; Maruthy et al., 2017), and if the layer V responses show passive currents from the other cortical layers or if they reflect active currents in the layer V outputs.

## REFERENCES

1. Abel TJ, Osorio RV, Amorim-Leite R, Mathieu F, Kahane P, Minotti L, Hoffmann D, Chabardes S (2018) Frameless robot-assisted stereoelectroencephalography in children: technical aspects and comparison with Talairach frame technique. Journal of Neurosurgery: Pediatrics 22:37–46.

2. Abrams DA, Kraus N (2005) Biomap: A neurodiognostic tool for auditory processing disorders. The ASHA Leader Available at: http://www.asha.org/Publications/leader/2005/051018/f051018c/ [Accessed January 22, 2016].

3. Anderson S, Parbery-Clark A, White-Schwoch T, Kraus N (2012) Aging Affects Neural Precision of Speech Encoding. Journal of Neuroscience 32:14156–14164.

4. Ayala YA, Lehmann A, Merchant H (2017) Monkeys share the neurophysiological basis for encoding sound periodicities captured by the frequency-following response with humans. Scientific Reports 7:1–11.

5. Bajo VM, Moore DR (2005) Descending projections from the auditory cortex to the inferior colliculus in the gerbil, Meriones unguiculatus. J Comp Neurol 486:101–116.

6. Bajo VM, Nodal FR, Moore DR, King AJ (2010) The descending corticocollicular pathway mediates learning-induced auditory plasticity. Nature Neuroscience 13:253–260.

7. Behroozmand R, Oya H, Nourski KV, Kawasaki H, Larson CR, Brugge JF, Howard MA, Greenlee JDW (2016) Neural Correlates of Vocal Production and Motor Control in Human Heschl’s Gyrus. J Neurosci 36:2302–2315.

8. Bidelman GM (2015) Multichannel recordings of the human brainstem frequency-following response: Scalp topography, source generators, and distinctions from the transient ABR. Hearing Research 323:68–80.

9. Bidelman GM (2018) Subcortical sources dominate the neuroelectric auditory frequency-following response to speech. NeuroImage 175:56–69.

10. Bidelman GM, Villafuerte JW, Moreno S, Alain C (2014) Age-related changes in the subcortical-cortical encoding and categorical perception of speech. Neurobiol Aging 35:2526–2540.

11. Chabardes S, Abel TJ, Cardinale F, Kahane P (2018) Commentary: Understanding Stereoelectroencephalography: What’s Next? Neurosurgery 82:E15–E16.

12. Chandrasekaran B, Hornickel J, Skoe E, Nicol T, Kraus N (2009) Context-Dependent Encoding in the Human Auditory Brainstem Relates to Hearing Speech in Noise: Implications for Developmental Dyslexia. Neuron 64:311–319.

13. Chandrasekaran B, Kraus N (2010a) The scalp-recorded brainstem response to speech: Neural origins and plasticity. Psychophysiology 47:236–246.

14. Chandrasekaran B, Kraus N (2010b) The scalp-recorded brainstem response to speech: Neural origins and plasticity. Psychophysiology 47:236–246.

15. Chou M-S, Lin C-D, Wang T-C, Jeng F-C (2014) Recording Frequency-following Responses to Voice Pitch in Guinea Pigs: Preliminary Results. Percept Mot Skills 118:681–690.

16. Coffey EBJ, Arseneau-Bruneau I, Zhang X, Baillet S, Zatorre RJ (2021) Oscillatory entrainment of the Frequency Following Response in auditory cortical and subcortical structures. J Neurosci Available at: https://www.jneurosci.org/content/early/2021/03/16/JNEUROSCI.2313-20.2021 [Accessed April 13, 2021].

17. Coffey EBJ, Chepesiuk AMP, Herholz SC, Baillet S, Zatorre RJ (2017a) Neural Correlates of Early Sound Encoding and their Relationship to Speech-in-Noise Perception. Front Neurosci 11 Available at: https://www.ncbi.nlm.nih.gov/pmc/articles/PMC5575455/ [Accessed March 30, 2018].

18. Coffey EBJ, Herholz SC, Chepesiuk AMP, Baillet S, Zatorre RJ (2016) Cortical contributions to the auditory frequency-following response revealed by MEG. Nat Commun 7:11070.

19. Coffey EBJ, Musacchia G, Zatorre RJ (2017b) Cortical Correlates of the Auditory Frequency-Following and Onset Responses: EEG and fMRI Evidence. The Journal of Neuroscience 37:830–838.

20. Coffey EBJ, Nicol T, White-Schwoch T, Chandrasekaran B, Krizman J, Skoe E, Zatorre RJ, Kraus N (2019) Evolving perspectives on the sources of the frequency-following response. Nat Commun 10:1–10.

21. Delorme A, Makeig S (2004) EEGLAB: an open source toolbox for analysis of single-trial EEG dynamics including independent component analysis. Journal of neuroscience methods 134:9–21.

22. Du Y, Ma T, Wang Q, Wu X, Li L (2009) Two crossed axonal projections contribute to binaural unmasking of frequency-following responses in rat inferior colliculus. European Journal of Neuroscience 30:1779–1789.

23. Faraji AH, Remick M, Abel TJ (2020) Contributions of Robotics to the Safety and Efficacy of Invasive Monitoring With Stereoelectroencephalography. Front Neurol 11 Available at: https://www.frontiersin.org/articles/10.3389/fneur.2020.570010/full [Accessed February 1, 2021].

24. Fischl B (2012) FreeSurfer. Neuroimage 62:774–781.

25. Galbraith GC, Bagasan B, Sulahian J (2001) Brainstem frequency-following response recorded from one vertical and three horizontal electrode derivations. Percept Mot Skills 92:99–106.

26. Galbraith GC, Threadgill MR, Hemsley J, Salour K, Songdej N, Ton J, Cheung L (2000) Putative measure of peripheral and brainstem frequency-following in humans. Neuroscience letters 292:123–127.

27. Gardi J, Merzenich M, McKean C (1979) Origins of the Scalp-Recorded Frequency-Following Response in the Cat. Audiology 18:353–380.

28. Gerken GM, Moushegian G, Stillman RD, Rupert AL (1975) Human frequency-following responses to monaural and binaural stimuli. Electroencephalogr Clin Neurophysiol 38:379–386.

29. Gnanateja GN, Ranjan R, Firdose H, Sinha SK, Maruthy S (2013) Acoustic basis of context dependent brainstem encoding of speech. Hearing Research 304:28–32.

30. Gnanateja GN, Ranjan R, Sandeep M (2012) Physiological bases of the encoding of speech evoked frequency following responses. Journal of All India Institute of Speech and Hearing 31:215–219.

31. Gorina-Careta N, Kurkela JLO, Hämäläinen J, Astikainen P, Escera C (2021) Neural generators of the frequency-following response elicited to stimuli of low and high frequency: A magnetoencephalographic (MEG) study. NeuroImage 231:117866.

32. Greenberg RP, Stablein DM, Becker DP (1981) Noninvasive localization of brain-stem lesions in the cat with multimodality evoked potentials: Correlation with human head-injury data. Journal of Neurosurgery 54:740–750.

33. Grimsley JMS, Shanbhag SJ, Palmer AR, Wallace MN (2012) Processing of Communication Calls in Guinea Pig Auditory Cortex. PLOS ONE 7:e51646.

34. Guo N, Si X, Zhang Y, Ding Y, Zhou W, Zhang D, Hong B (2020) Speech frequency-following response in human auditory cortex is more than a simple tracking. NeuroImage:117545.

35. Hartmann T, Weisz N (2019) Auditory cortical generators of the Frequency Following Response are modulated by intermodal attention. NeuroImage 203:116185.

36. He W, Ding X, Zhang R, Chen J, Zhang D, Wu X (2014) Electrically-Evoked Frequency-Following Response (EFFR) in the Auditory Brainstem of Guinea Pigs. PLOS ONE 9:e106719.

37. Heffner HE, Heffner RS (2007) Hearing Ranges of Laboratory Animals. Journal of the American Association for Laboratory Animal Science 46:20–22.

38. Kaas JH, Hackett TA (2000) Subdivisions of auditory cortex and processing streams in primates. PNAS 97:11793–11799.

39. King A, Hopkins K, Plack CJ (2016) Differential Group Delay of the Frequency Following Response Measured Vertically and Horizontally. JARO 17:133–143.

40. Kraus N, Thompson EC, Krizman J, Cook K, White-Schwoch T, LaBella CR (2016) Auditory biological marker of concussion in children. Sci Rep 6 Available at: https://www.ncbi.nlm.nih.gov/pmc/articles/PMC5178332/ [Accessed June 26, 2020].

41. Krishnan A, Bidelman GM, Gandour JT (2010a) Neural representation of pitch salience in the human brainstem revealed by psychophysical and electrophysiological indices. Hearing Research 268:60–66.

42. Krishnan A, Gandour JT, Bidelman GM (2010b) The effects of tone language experience on pitch processing in the brainstem. Journal of Neurolinguistics 23:81–95.

43. Krishnan A, Gandour JT, Bidelman GM (2012) Experience-dependent plasticity in pitch encoding: from brainstem to auditory cortex. NeuroReport 23:498–502.

44. Krizman J, Kraus N (2019) Analyzing the FFR: A tutorial for decoding the richness of auditory function. Hearing Research 382:107779.

45. Lau JCY, Wong PCM, Chandrasekaran B (2017) Context-dependent plasticity in the subcortical encoding of linguistic pitch patterns. J Neurophysiol 117:594–603.

46. Lau JCY, Wong PCM, Chandrasekaran B (2018) Interactive effects of linguistic abstraction and stimulus statistics in the online modulation of neural speech encoding. Atten Percept Psychophys Available at: https://doi.org/10.3758/s13414-018-1621-9 [Accessed April 18, 2019].

47. Li F, Teichert T (2020) A surface metric and software toolbox for EEG electrode grids in the macaque. Journal of Neuroscience Methods 346:108906.

48. Liégeois-Chauvel C, Musolino A, Badier JM, Marquis P, Chauvel P (1994) Evoked potentials recorded from the auditory cortex in man: evaluation and topography of the middle latency components. Electroencephalography and Clinical Neurophysiology/Evoked Potentials Section 92:204–214.

49. Llanos F, Xie Z, Chandrasekaran B (2017) Hidden Markov modeling of frequency-following responses to Mandarin lexical tones. Journal of Neuroscience Methods 291:101–112.

50. Makeig S, Debener S, Onton J, Delorme A (2004) Mining event-related brain dynamics. Trends in Cognitive Sciences 8:204–210.

51. Marsh JT, Brown WS, Smith JC (1974) Differential brainstem pathways for the conduction of auditory frequency-following responses. Electroencephalogr Clin Neurophysiol 36:415–424.

52. Marsh JT, Brown WS, Smith JC (1975) Far-field recorded frequency-following responses: correlates of low pitch auditory perception in humans. Electroencephalogr Clin Neurophysiol 38:113–119.

53. Maruthy S, Kumar UA, Gnanateja GN (2017) Functional Interplay Between the Putative Measures of Rostral and Caudal Efferent Regulation of Speech Perception in Noise. Journal of the Association for Research in Otolaryngology 18:635–648.

54. Naert G, Pasdelou M-P, Le Prell CG (2019) Use of the guinea pig in studies on the development and prevention of acquired sensorineural hearing loss, with an emphasis on noise. The Journal of the Acoustical Society of America 146:3743–3769.

55. Parbery-Clark A, Strait DL, Kraus N (2011) Context-dependent encoding in the auditory brainstem subserves enhanced speech-in-noise perception in musicians. Neuropsychologia 49:3338–3345.

56. Peng F, Innes-Brown H, McKay CM, Fallon JB, Zhou Y, Wang X, Hu N, Hou W (2018) Temporal Coding of Voice Pitch Contours in Mandarin Tones. Front Neural Circuits 12 Available at: https://www.frontiersin.org/articles/10.3389/fncir.2018.00055/full [Accessed April 27, 2021].

57. Petkov CI, Kayser C, Augath M, Logothetis NK (2006) Functional Imaging Reveals Numerous Fields in the Monkey Auditory Cortex. PLOS Biology 4:e215.

58. Plack CJ, Barker D, Hall DA (2014) Pitch coding and pitch processing in the human brain. Hearing Research 307:53–64.

59. Plyler PN, Ananthanarayan AK (2001) Human frequency-following responses: representation of second formant transitions in normal-hearing and hearing-impaired listeners. J Am Acad Audiol 12:523–533.

60. Presacco A, Simon JZ, Anderson S (2016) Evidence of degraded representation of speech in noise, in the aging midbrain and cortex. Journal of Neurophysiology 116:2346–2355.

61. Rauschecker JP, Tian B (2000) Mechanisms and streams for processing of “what” and “where” in auditory cortex. PNAS 97:11800–11806.

62. Reetzke R, Xie Z, Llanos F, Chandrasekaran B (2018) Tracing the Trajectory of Sensory Plasticity across Different Stages of Speech Learning in Adulthood. Current Biology 28:1419–1427.e4.

63. Sinnott JM, Beecher MD, Moody DB, Stebbins WC (1976) Speech sound discrimination by monkeys and humans. The Journal of the Acoustical Society of America 60:687–695.

64. Sinnott JM, Kreiter NA (1991) Differential sensitivity to vowel continua in Old World monkeys (Macaca) and humans. The Journal of the Acoustical Society of America 89:2421–2429.

65. Skoe E, Kraus N (2010a) Auditory brain stem response to complex sounds: a tutorial. Ear Hear 31:302–324.

66. Skoe E, Kraus N (2010b) Hearing It Again and Again: On-Line Subcortical Plasticity in Humans Op de Beeck HP, ed. PLoS ONE 5:e13645.

67. Slabu L, Grimm S, Escera C (2012) Novelty Detection in the Human Auditory Brainstem. J Neurosci 32:1447–1452.

68. Smith JC, Marsh JT, Brown WS (1975) Far-field recorded frequency-following responses: evidence for the locus of brainstem sources. Electroencephalogr Clin Neurophysiol 39:465–472.

69. Smith JC, Marsh JT, Greenberg S, Brown WS (1978) Human auditory frequency-following responses to a missing fundamental. Science 201:639–641.

70. Steinschneider M, Fishman YI, Arezzo JC (2003) Representation of the voice onset time (VOT) speech parameter in population responses within primary auditory cortex of the awake monkey. The Journal of the Acoustical Society of America 114:307–321.

71. Steinschneider M, Volkov IO, Noh MD, Garell PC, Howard MA (1999) Temporal Encoding of the Voice Onset Time Phonetic Parameter by Field Potentials Recorded Directly From Human Auditory Cortex. Journal of Neurophysiology 82:2346–2357.

72. Tadel F, Baillet S, Mosher JC, Pantazis D, Leahy RM (2011) Brainstorm: A User-Friendly Application for MEG/EEG Analysis. Available at: https://www.hindawi.com/journals/cin/2011/879716/ [Accessed June 4, 2018].

73. Teichert T (2016) Tonal frequency affects amplitude but not topography of rhesus monkey cranial EEG components. Hearing Research 336:29–43.

74. Teichert T, Gnanateja GN, Sadagopan S, Chandrasekaran B (2021) Linear superposition of responses evoked by individual glottal pulses explain over 80% of the frequency following response to human speech in the macaque monkey. bioRxiv:2021.09.06.459204.

75. Teichert T, Gurnsey K, Salisbury D, Sweet RA (2016) Contextual processing in unpredictable auditory environments: the limited resource model of auditory refractoriness in the rhesus. Journal of Neurophysiology 116:2125–2139.

76. Vander Werff KR, Rieger B (2017) Brainstem Evoked Potential Indices of Subcortical Auditory Processing Following Mild Traumatic Brain Injury. Ear Hear 38:e200–e214.

77. Wallace MN, Rutkowski RG, Palmer AR (2000) Identification and localisation of auditory areas in guinea pig cortex. Exp Brain Res 132:445–456.

78. Wang Q, Li L (2018) Differences between auditory frequency-following responses and onset responses: Intracranial evidence from rat inferior colliculus. Hearing Research 357:25–32.

79. White-Schwoch T, Carr KW, Thompson EC, Anderson S, Nicol T, Bradlow AR, Zecker SG, Kraus N (2015) Auditory Processing in Noise: A Preschool Biomarker for Literacy. PLOS Biology 13:e1002196.

80. Worden FG, Marsh JT (1968) Frequency-following (microphonic-like) neural responses evoked by sound. Electroencephalogr Clin Neurophysiol 25:42–52.

81. Yamada O, Marsh RR, Potsic WP (1980) Generators of the Frequency-following Response in the Guinea Pig. Otolaryngol Head Neck Surg 88:613–618.

82. Zatorre RJ, Belin P (2001) Spectral and Temporal Processing in Human Auditory Cortex. Cereb Cortex 11:946–953.

